# Vinculin controls endothelial cell junction dynamics during vascular lumen formation

**DOI:** 10.1101/2021.03.08.434441

**Authors:** Maria P. Kotini, Miesje M. van der Stoel, Mitchell K. Han, Bettina Kirchmaier, Johan de Rooij, Markus Affolter, Stephan Huveneers, Heinz-Georg Belting

## Abstract

Blood vessel morphogenesis is driven by coordinated endothelial cell behaviors, which depend on dynamic cell-cell interactions. Remodeling of endothelial cell-cell junctions promote morphogenetic cellular events while preserving vascular integrity. Here, we have analyzed the dynamics of endothelial cell-cell junctions during lumen formation in angiogenic sprouts. By live-imaging of the formation of intersegmental blood vessels in zebrafish, we demonstrate that lumen expansion is accompanied by the formation of transient finger-shaped junctions. Formation and maintenance of these junctional fingers are positively regulated by blood pressure whereas inhibition of blood flow prevents their formation. Using fluorescent reporters, we show that the tension-sensor Vinculin localizes to junctional fingers. Furthermore, loss of vinculin function, in *vinculin a* and -*b* double knockouts, prevents junctional finger formation in angiogenic sprouts, whereas endothelial expression of a *vinculin* transgene is sufficient to restore junctional fingers. Taken together, our findings suggest a mechanism in which lumen expansion during angiogenesis leads to an increase in junctional tension, which triggers recruitment of vinculin and formation of junctional fingers. We propose that endothelial cells may employ force-dependent junctional remodeling to react to changes in external forces to protect cell-cell contacts and to maintain vascular integrity during sprouting angiogenesis.

## Introduction

The cardiovascular system is essential for vertebrate development and homeostasis as it serves as a transport system to deliver and exchange gas and solutes in all parts of the body. The cardiovascular system is the first organ to form during embryonic development and it adapts to the needs of the growing embryo by expansion of the vascular network (reviewed by Hogan and Schulte-Merker, 2017; Potente et al., 2011). This is to a large extent achieved by the formation of new vessels in a process called sprouting angiogenesis.

Blood vessel formation by angiogenesis occurs in discrete steps of sprout outgrowth, lumen formation and sprout or vessel fusion. At the cellular level, these morphogenetic processes are achieved by numerous cell activities such as endothelial cell migration, cell rearrangements and cell shape changes, which in turn are driven by a complex array of intracellular activities as well as dynamic cell-cell and cell-ECM interactions (reviewed by Betz et al., 2016; Szymborska and Gerhardt, 2018). During angiogenesis, multiple morphogenetic processes are occurring simultaneously, posing a particular challenge to endothelial cell-cell junctions. Fine-tuned regulation of cell-cell adhesion is required to permit endothelial cell migration and rearrangements while maintaining the vascular seal to prevent leakage or hemorrhage.

Endothelial cell-cell junctions play a central role in many aspects of endothelial cell biology. They regulate not only cell-cell adhesion and vascular permeability, but also proliferation, survival, shape and motility, and they are involved in the modulation of signaling pathways and transcriptional regulation (reviewed by Komarova et al., 2017; Lampugnani et al., 2017; Wallez and Huber, 2008). In addition to forming a structural link between neighboring endothelial cells, the junctions can act as sensors to adapt to forces that are derived from the actomyosin cytoskeleton or the vascular hemodynamic microenvironment (Huveneers et al., 2012; Lagendijk et al., 2017; Tzima et al., 2005; Liu et al., 2010). During vascular morphogenesis, endothelial cells are exposed to mechanical forces generated by matrix stiffness, fluid shear stress, blood pressure, actomyosin contractility and junctional tension (Duchemin et al., 2019; Hoefer et al., 2013; Roman and Pekkan, 2012). Recent work has shown that endothelial cell-cell junctions play an important role in mechanical force sensing and force transmission, specifically by changing the organization of VE-cadherin-based junctions (Angulo-Urarte et al., 2020; Baeyens and Schwartz, 2016; Conway et al., 2013). Tension on the vascular endothelial (VE)-cadherin/β-catenin/α-catenin complex recruits the mechanotransduction protein Vinculin, which in turn protects the endothelial junctions during their remodeling (Huveneers et al., 2012). Force-dependent signals are transmitted through the VE-cadherin complex and via junctional Vinculin recruitment enable collective endothelial responses *in vitro* (Barry et al., 2015).

Recent work in zebrafish embryos showed that the adhesion molecule VE-cadherin transduces intracellular actomyosin contractility to endothelial cell-cell junctions *in vivo* (Lagendijk et al., 2017) and also participates in force generating processes during endothelial cell rearrangements (Paatero et al., 2018). Here, we have studied how endothelial cells respond to mechanical stress imposed in a changing mechanical environment. To this end we examine endothelial cell-cell junctions during angiogenic lumen expansion, which is accompanied with an acute increase in blood pressure within the sprouting intersegmental vessels (ISVs). Our studies reveal junctional structures within angiogenic sprouts, called junctional fingers, which form transiently in response to increased pressure and which recruit Vinculin. Genetic analyses reveal that Vinculin is required for junctional finger formation. Taken together, our studies indicate that endothelial cells protect their cell-cell contact against mechanical stress by forming junctional fingers, thereby maintaining sprout integrity during lumen expansion.

## Results

### Endothelial junctional fingers appear during vascular lumen formation in vivo

During blood vessel formation endothelial junctions play permissive as well as active roles, driving cell rearrangements and cell elongation while maintaining vascular integrity at the same time (Paatero et al., 2018; Sauteur et al., 2014; reviewed by Fonseca et al., 2020; Okuda and Hogan, 2020). Furthermore, endothelial junctions are thought to sense and transmit mechanical forces and thus regulate the collective endothelial response to changes in blood flow or pressure, which occur at the onset of blood flow in a newly formed vessel. To investigate the response of endothelial cell-cell junctions to changes in blood pressure, we followed junctional dynamics in zebrafish ISVs during the onset of lumen formation around 28-36 hours post fertilization (hpf). Immunofluorescent analysis of the adherens junction protein VE-cadherin (Cdh5) and tight junction protein zonula occludens-1 (ZO-1) revealed finger-shaped structures, which emanated from the cell-cell contact interface, confirming our previous observations (Sauteur et al., 2014). These junctional fingers appeared perpendicular to the originating junction with an angle of 85-95 degrees (Fig. 1 A). To check whether these junctional fingers are specifically formed by VE-cadherin-based adhesions or represent more general junctional structures, we analyzed the localization of platelet endothelial cell adhesion molecule-1 (Pecam1) using a transgenic reporter *(Tg(fli1a:pecam1-EGFP)^ncv27^*), followed by immunofluorescent analysis of ZO-1 (Fig. 1 B). We found that junctional fingers also contained Pecam1 and colocalized with ZO-1, indicating that junctional fingers were formed by several types of cell-cell adhesion complexes.

**Figure 1.**
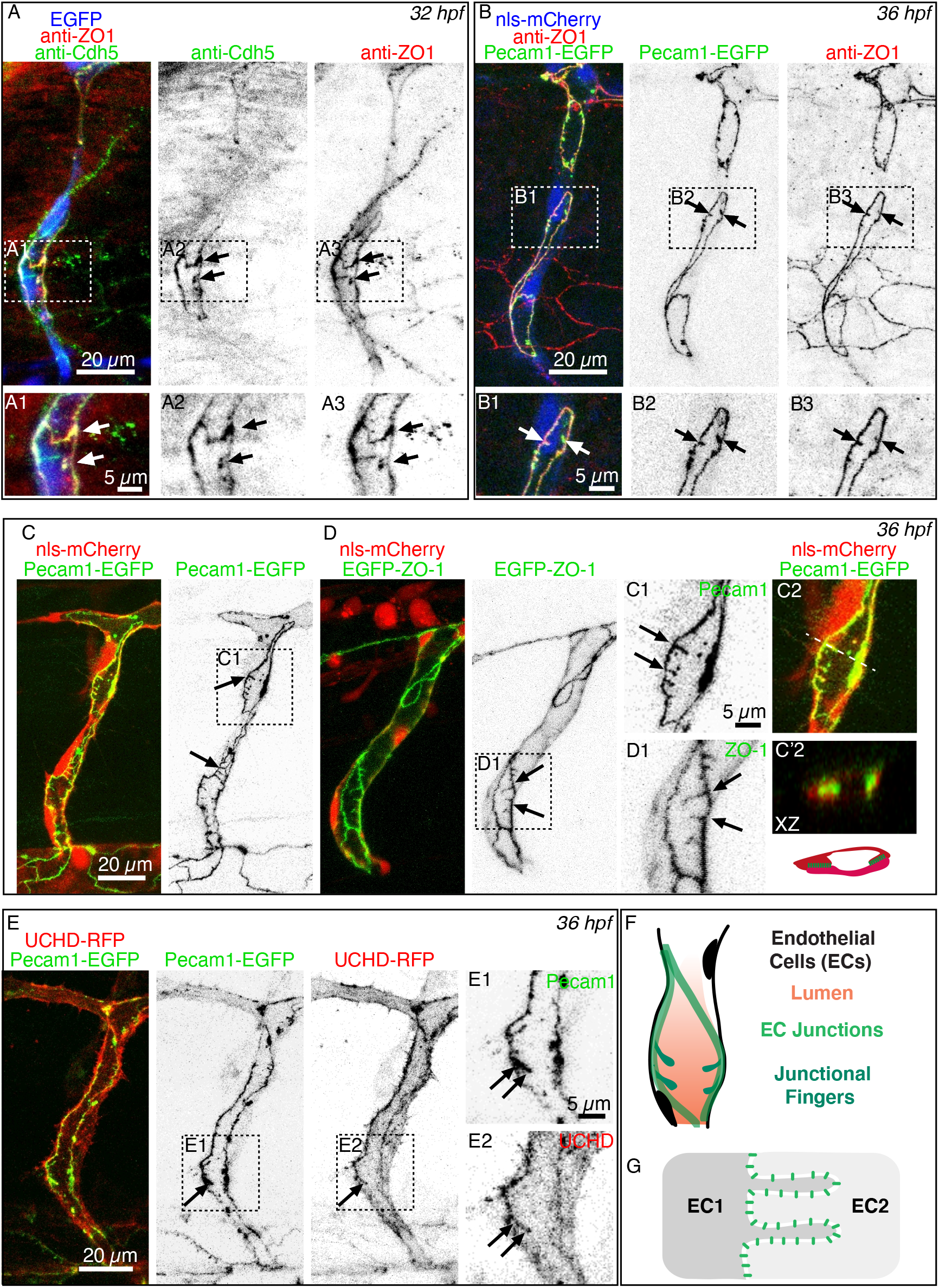
Endothelial junction fingers are formed by distinct junctional proteins. Confocal images of intersegmental blood vessels (ISVs) in the trunk of fixed zebrafish embryos at 32 hpf (A) and 36 hpf (B - E). **(A)** Whole mount immunofluorescence staining using anti-Cdh5 (VE-cadherin) and anti-ZO-1 in *Tg(fli:EGFP)* embryos. Scale bars, 20 μm. **(A1-3)** Zoom-in images of the outlined boxes showing junctional fingers containing VE-cadherin and ZO-1. Scale bars, 5 μm. **(B)** Whole mount immunofluorescence staining using anti-ZO-1 in *Tg(fli:nls-mCherry;fli:pecam1-EGFP)* embryos. Scale bars, 20 μm. **(B1-3)** Zoom-in images of the outlined boxes in B, showing the presence of PECAM-1 and ZO-1 at junctional fingers between endothelial cells. Scale bars, 5 μm. **(C)** GFP+ signal shows the junctional molecule Pecam1 and RFP+ signal marks endothelial cell nuclei in *Tg(fli:nls-mCherry;fli:pecam1-EGFP)* embryos. Scale bar, 20 μm. **(D)** GFP+ signal marks the junctional molecule ZO-1 and RFP+ signal indicates endothelial cell nuclei in *Tg(fli:nls-mCherry;fli:gal4ff;UAS:EGFP-ZO-1)* embryos. Scale bar, 20 μm. **(C1, D1)** Zoom-in images of the outlined boxes in C - D, black arrows indicate the presence of Pecam1 and ZO-1 at junctional fingers. Scale bars, 5 μm. **(C2-C’2)** XZ representation of junctional finger in C2. **(E)** GFP+ signal shows the junctional molecule Pecam1 and RFP+ signal marks the F-actin cytoskeleton in *Tg(fli:gal4ff;4xUAS:UCHD-RFP;fli:pecam1-EGFP)* embryos. Scale bars, 20 μm. **(E1-2)** Zoom-in images of the outlined boxes in E. Black arrow indicates the presence of F-actin at a junctional finger. Scale bars, 5 μm. **(F)** Schematic representation of junctional fingers at the interface of two endothelial cells. (G) Schematic representation of junctional fingers appearing in an ISV during lumen formation.

Because fixation of zebrafish embryos disrupts blood flow and can affect blood vessel morphology, we wanted to verify the existence of junctional fingers by live-imaging using the above-mentioned Pecam1 reporter (Fig. 1 C) as well as a ZO-1 reporter *Tg(UAS:EGFP-hZO1)^ubs5^* (Fig. 1 D). These experiments confirmed the presence of ZO-1 and Pecam1-positive fingers in the lumenized ISVs (Fig. 1 C-D). Furthermore, junctional fingers in live embryos appeared more prominent and larger compared to those seen in fixed embryos, suggesting that they are sensitive to the loss of blood pressure or fixation. Because cell-cell junctions are physically connected to the actin cytoskeleton, we next tested whether junctional fingers colocalize with Ruby-UCHD, which labels stabilized F-actin fibers. Live imaging revealed that Pecam1-positive junctional fingers were also marked by mRuby2-UCHD (Fig. 1 E). Taken together, this set of experiments indicate that junctional fingers are formed during angiogenic sprouting in the developing ISVs and contain multiple junctional proteins, including VE-cadherin, ZO-1 and Pecam1, which can interact with the endothelial F-actin cytoskeleton.

Since junctional fingers are particularly prevalent in newly lumenized blood vessels (30-36 hpf), we monitored the appearance of junctional fingers during this time window by *in vivo* time-lapse imaging using the Pecam1-EGFP reporter to investigate what controls their formation. Before ISV lumen opening, we observed the presence of dense junctional clusters, but no junctional fingers (Fig. 2, time points 264 and 384 min, and Supplementary Movie 1). By contrast, upon lumen opening the junctional fingers readily formed (Fig. 2; from time point 408 min onwards and Supplementary Movie 1). Over time, the size of the fingers fluctuated and they eventually regressed once the ISV lumens are connected upon anastomosis, and mostly stable, linear-oriented junctions are established (Fig. 2 B-C; time points: 432, 456, and 528 min, red arrows and Supplementary Movie 1). These findings show that junctional fingers are transient structures, which form during lumen formation prior to commencement of blood flow. When blood flow is established the junctional fingers disappear. These correlations suggest a close relationship between hemodynamic forces and junctional finger formation.

**Figure 2.**
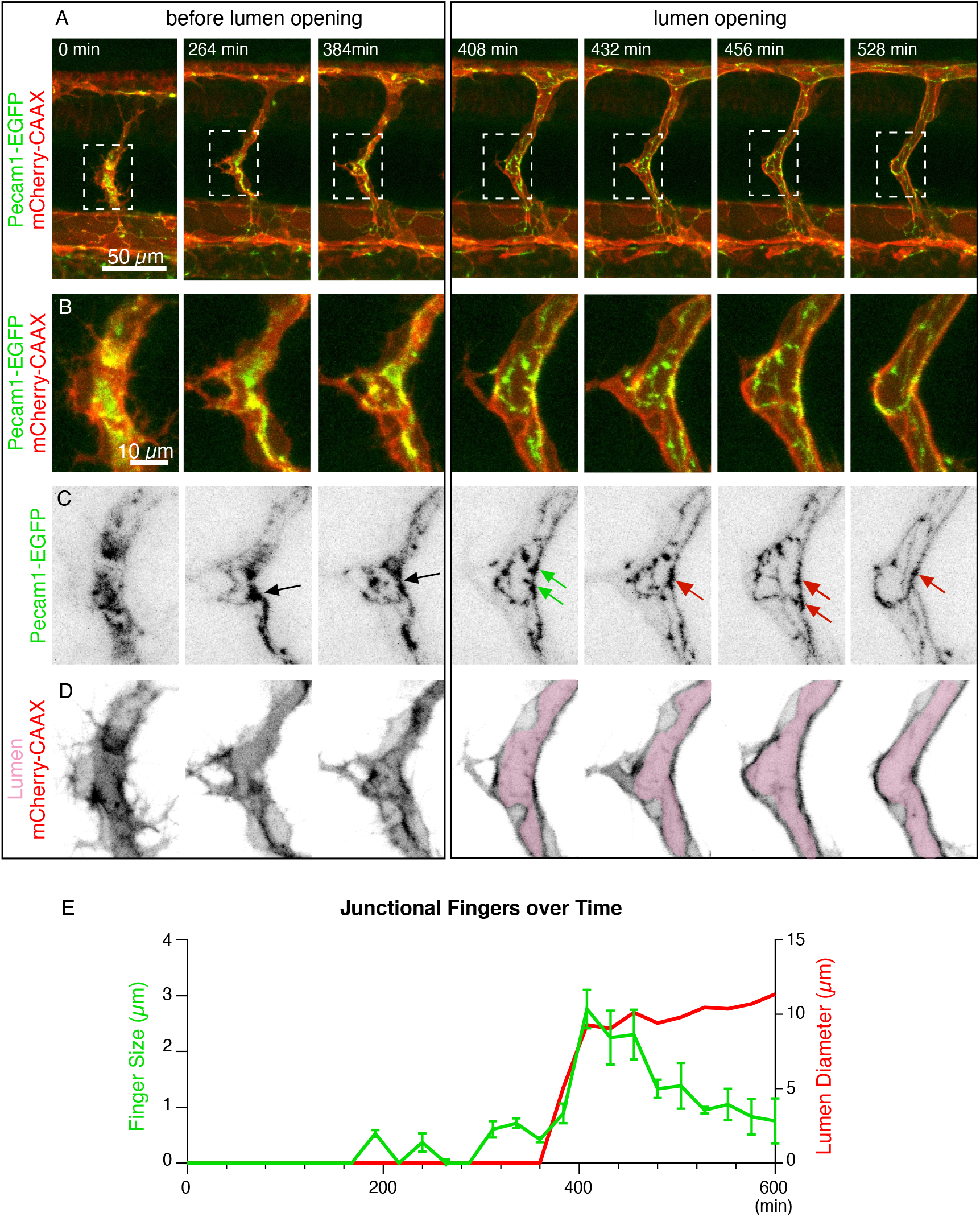
Junctional fingers assemble upon lumen opening. **(A-D)** Still images from a time lapse movie of a *Tg(kdrl:mCherry-CAAX;fli:pecam1-EGFP)* embryo starting at 32 hpf. GFP+ signal marks the endothelial cell-cell junctions and mCherry+ signal demarcates endothelial cell plasma membranes. First three time points (0, 264, 384 min) display a representative ISV before lumen opening. The following time points (408; 432; 456; 528 min) visualize the ISV after lumen opening. **(A)** Overview images. Scale bars, 50 μm. **(B)** Zoom-in images of the outlined boxes in A. Scale bars, 10 μm. **(C)** Inverted contrast of the Pecam1-EGFP channel from B. Right before lumen opening, junctional accumulation is evident (black arrows) at future sites of junctional fingers. After lumen opening, junctional fingers assemble (green arrows) and disassemble (red arrows) over time. **(D)** Inverted contrast of mCherry-CAAX from B showing lumen opening. Pink pseudo-color indicates the vascular lumen. Scale bars (C-D), 10 μm. **(E)** Line graph analysis showing the junctional finger size ±S.D. and lumen diameter over time (0 to 600 min) from a representative *Tg(kdrl:mCherry-CAAX;fli:pecam1-EGFP)* embryo. **See also Supplementary Movie 1.**

### Hemodynamic forces control junctional fingers

Previous studies have shown that ISVs can form lumens in different ways. Cell membrane invagination leads to the formation of a lumen within a cell (Herwig et al., 2011). This process, also called transcellular lumen formation, is driven by blood pressure and thus requires blood flow (Gebala et al., 2016; Herwig et al., 2011; Lenard et al., 2013). In the absence of blood flow ISV lumens are formed by cord hollowing, which is driven by endothelial cell rearrangements and gives rises to multicellular tubes (Herwig et al, 2011). Because ISV sprouting is initiated just prior to the start of embryonic blood flow, both modes of lumen formation may occur (Gebala et al., 2016; Herwig et al., 2011).

Because of the correlation between the occurrence of junctional fingers and lumen formation on one hand and the relationship between blood flow and lumen inflation on the other, we wanted to address to what extent the formation of junctional fingers is dependent on hemodynamic forces. We applied different drug treatments to modulate blood flow in wild-type embryos with transgenic endothelial reporters for cell-cell junctions *(fli:pecam1-EGFP)* and cell membranes *(fli:mCherry-CAAX).* Tricaine treatment, which reduces cardiac contraction and consequently inhibits blood flow and blood pressure (Lagendijk et al., 2017; Lenard et al., 2013), led to a decrease in size and number of fingers compared to controls (vehicle treated embryos) (Fig. 3). By contrast, norepinephrine (NE) treatment, which raises the heart rate and increases the hemodynamic forces in zebrafish blood vessels (Chen et al., 2012; De Luca et al., 2014) increased the number and length of junctional fingers (Fig. 3).

**Figure 3.**
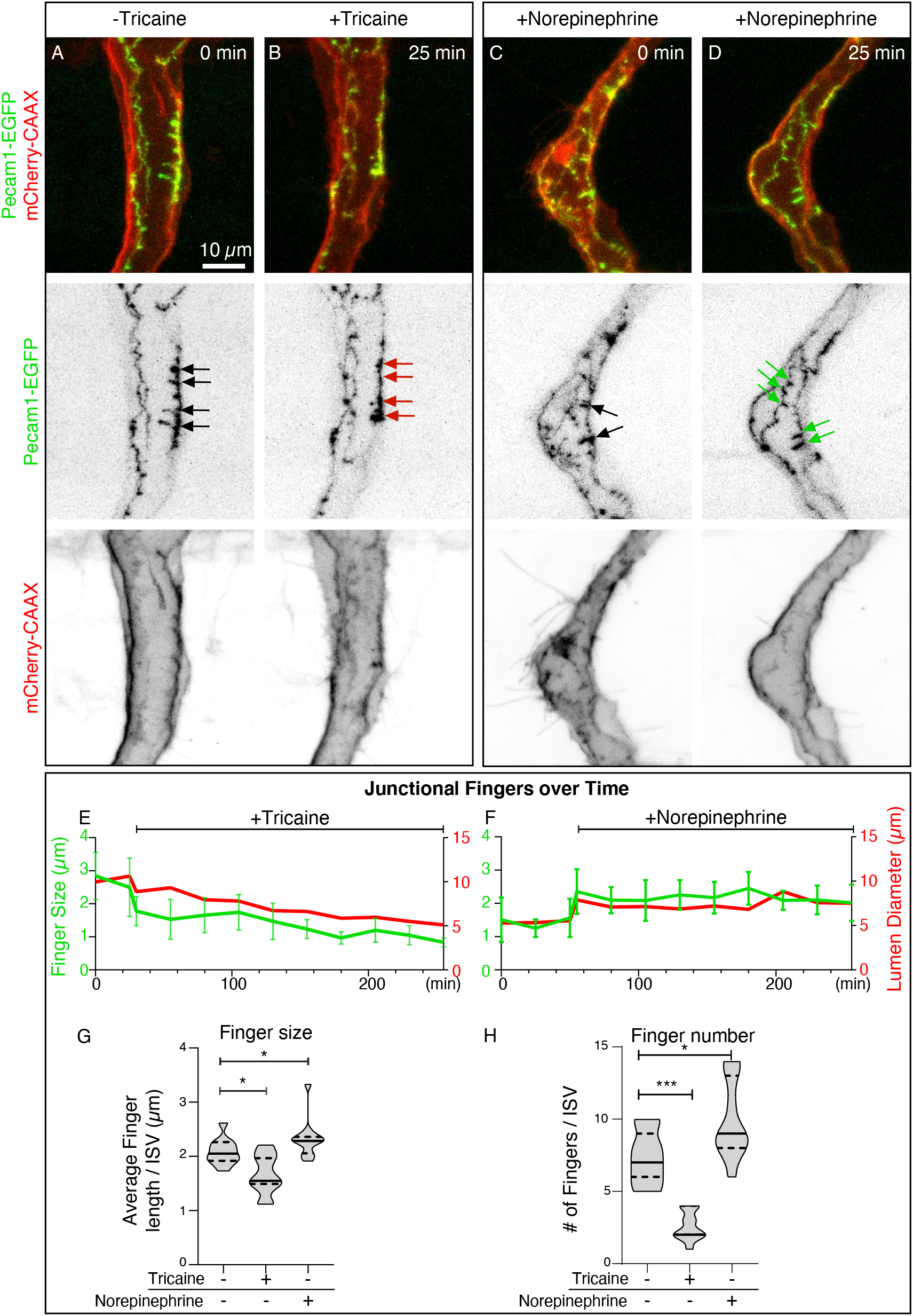
Alterations in blood flow affect finger length and number. Confocal images of representative ISVs from *Tg(kdrl:mCherry-CAAX;fli:pecam1-EGFP)* embryos at 36 hpf before and after treatment with 0.32% tricaine or 60 μm norepinephrine. GFP+ signal marks Pecam1 and RFP+ signal marks the cell membrane. **(A-D)** Merged and inverted contrast images of close ups of a representative ISV at 36 hpf, control before tricaine treatment **(A)**, after 25 min tricaine treatment **(B)**, control before norepinephrine treatment **(C)** and after 25 min of norepinephrine treatment **(D)**. Before pharmacological additions, junctional fingers are evident (A, C; black arrows) and upon tricaine treatment they regress (B; red arrows), while upon norepinphirine treatment they are reinforced (D; green arrows). Scale bar, 10 μm. **(E-F)** Line graphs showing the junctional finger size ±S.D. and lumen diameter over time after treatment with (E) tricaine or (F) norepinephrine from a representative *Tg(kdrl:mCherry-CAAX;fli:pecam1-EGFP)* embryo. **(G)** Violin plot showing the average finger length per ISV in non-treated, tricaine or norepinephrine treated embryos, *p<0.05 (One-Way ANOVA with Dunnett’s post-test, n = 11 ISVs from 8 embryos). **(H)** Violin plot showing the number of fingers per ISV in non-treated, tricaine or norepinephrine treated embryos, *p<0.05, ***p<0.001 (One-Way ANOVA with Dunnett’s post-test, n = 11 ISVs from 8 embryos).

To investigate whether changes in hemodynamic forces directly affect junctional finger dynamics, we next performed sequential treatments. Junctional clusters appeared before lumen opening of the ISV in control (*vcla*^+/-^;*vclb*^+/+^) zebrafish embryos (Fig. 4 A; time point 0; Supplementary Movie 2) and junctional fingers were formed during ISV lumenization (Fig. 4 A, time points 20, 60 min), confirming our earlier findings (Fig. 3). Subsequent treatment of the same embryo with NE to promote blood flow, induced junctional clustering and finger formation in the lumenized ISV (Fig. 4 B; time points: 120; 160 min after NE). Follow-up treatment with tricaine immediately caused lumen regression and destabilized junctional fingers (Fig. 4 C). Administration of the treatments in reversed order showed consistent results (Supplementary Fig. 1, Supplementary Movie 3). Taken together, these experiments show that junctional fingers form in response to an increase in blood flow, or pressure, and that they are dynamic structures that directly respond to changes in hemodynamic forces.

**Figure 4.**
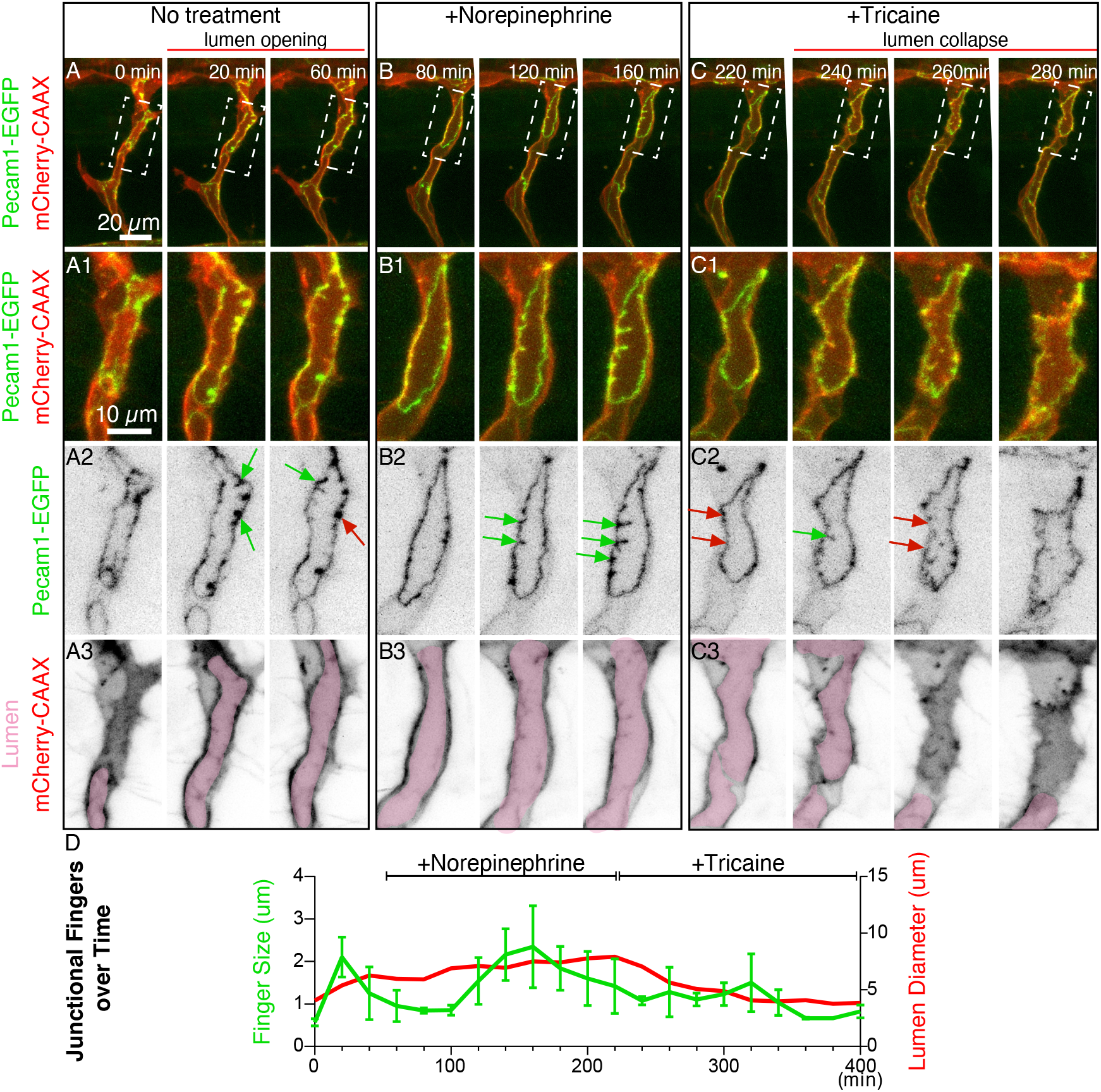
Hemodynamic forces control junctional fingers. **(A-C)** Stills of time lapse confocal imaging from a representative 36 hpf *Tg(kdrl:mCherry-CAAX;fli:pecam1-EGFP)* embryo before treatment (A) and after sequential additions of, first norepinephrine (B) and then of tricaine (C). GFP+ signal indicates the junctional marker Pecam-1 and RFP+ signal marks the endothelial cell membrane. Scale bars, 20 μm. **(A1-C1)** Zoom-in images of the outlined boxes in A-C. Scale bars, 10 μm. **(A2-C2)** Inverted contrast images of *pecam1-EGFP* signal of A1-C1. (A2) Before lumen opening, no junctional fingers are present, after lumen opening, Pecam1-based junctional fingers assemble (green arrows) and overtime undergo regression (red arrows). (B2) Upon norepinephrine treatment, Pecam1 assembles into enlarged junctional fingers (green arrows). (C2) After tricaine treatment, remaining Pecam1-EGFP positive fingers (green arrows) disappear over time (red arrows). **(A3-C3)** Inverted contrast images of mCherry-CAAX of A1-C1. Pink pseudo-colored areas indicate open lumen in the ISV. (A3) Lumen opens up. (B3) After norepinephrine addition lumen opening expands. (C3) Treatment with tricaine results in lumen compartmentalization and lumen regression. **(D)** Line graph analysis showing the junctional finger length ±S.D. and lumen diameter over time before treatment (0 - 40 minutes), after treatment with norepinephrine (60 to 220 minutes) and after treatment with tricaine (240 to 400 minutes) from the same *Tg(kdrl:mCherry-CAAX;fli: pecam1-EGFP)* embryo. See also supplementary movie 2.

### Junctional fingers represent regions of elevated junctional tension

The observation that junctional fingers form in response to increased blood pressure suggests that their formation may be triggered by mechanical forces. We therefore reasoned that an acute increase in blood pressure may increase tensile forces exerted on endothelial cell-cell junctions. Previous work in HUVECs has shown that increased tension on the VE-cadherin complex induces unfolding of α-catenin, exposing its cryptic Vinculin binding site (Huveneers et al., 2012; Yao et al., 2014). The subsequent junctional recruitment of Vinculin reinforces the connection of VE-cadherin with the actin cytoskeleton and protects junctions during their force-dependent remodeling (Huveneers et al., 2012). Thus, since junctional localization of Vinculin corresponds to sites of increased tension, we wanted to test whether Vinculin is also localized at junctional fingers in ISVs. To achieve this, we transiently expressed a mCherry-Vinculin fusion protein *(kdrl:mCherry-vinculin)* in transgenic *flił:pecamł-EGFP* embryos. Using live imaging, we observed that Vinculin localizes at the junctional fingers (black arrows) (Fig. 5 A-C). Moreover, time-lapse imaging showed that Vinculin localization is dynamic and follows the turnover of the Pecam1-positive junctional fingers. Although the majority of Vinculin was distributed in the cytoplasm, upon formation of junctional fingers, Vinculin became enriched at these structures and was most prominent once junctional fingers reached their maximum length (Fig. 5 A-C, Supplementary Movie 4). This close correlation between junctional finger formation and Vinculin accumulation indicates that junctional fingers represent regions of elevated tension at the junctional interface. Furthermore, these observations suggest that Vinculin may play a role in the formation and/or function of junctional fingers.

**Figure 5.**
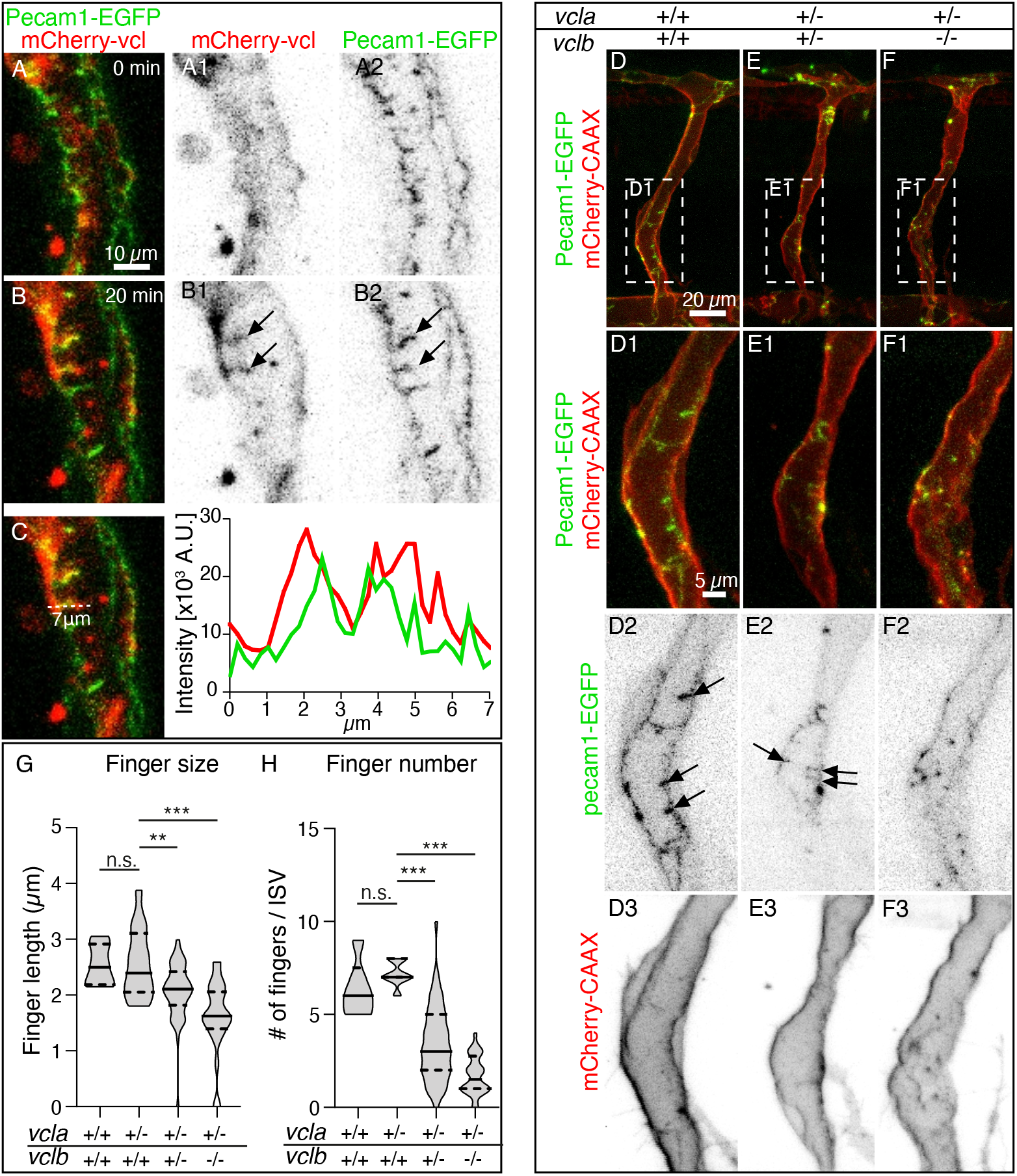
Vinculin is required for the formation of junctional fingers. **(A-B)** Stills of time lapse imaging of *Tg(fli:pecam1-EGFP)* embryos at 36 hpf with transient expression of injected *kdrl:mCherry-vinculin (vcl)* (Supplementary movie 4). Scale bar, 10 μm. **(A1-B1)** Inverted contrast images of mCherry-Vinculin of A-B. Red arrows point to colocalization of Vinculin and Pecam1. **(A2-B2)** Inverted contrast images of *fli:pecam1-EGFP* of A-B. Red arrows piont to co-localization of Vinculin and Pecam1 at junctional fingers. **(C)** Line scan analysis of a junctional finger showing the association of *kdrl:mCherry-vcl* and *fli:pecam1-EGFP* intensities. **See also supplementary movie 3. (D - F)** Confocal images of ISVs from (D) wild-type (*vcla*^+/+^;*vclb*^+/+^), (E) double heterozygous (*vcla*^+/-^;*vclb*^+/-^) or (F) *vclb* homozygous (*vcla*^+/-^;*vclb*^-/-^) *Tg(kdrl:mCherry-CAAX;fli:pecam1-EGFP)* embryos at 36 hpf. Scale bar, 20 μm. **(D1-F1)** Zoom-in images of outlined boxes in D-F. Scale bar, 5 μm. **(D2-F2)** Inverted contrast images *fli:pecam1-EGFP* of D1-F1. **(D3-F3)** Inverted contrast images of *mCherry-CAAX* of D1-F1. Black arrows indicate junctional fingers, note the absence of junctional fingers in *vclb* homozygous knock outs. **(G)** Violin plot showing the average finger length per ISV of wild-type (*vcla*^+/+^;*vclb*^+/+^), single heterozygous (*vcla*^+/-^;*vclb*^+/+^), double heterozygous (*vcla*^+/-^;*vclb*^+/-^) or *vclb* homozygous knockout (*vcla*^+/-^;*vclb*^-/-^) embryos (n = 4 (*vcla*^+/+^;*vclb*^+/+^), 9 (*vcla*^+/-^;*vclb*^+/+^), 22 (*vcla*^+/-^;*vclb*^+/-^) and 12 (*vcla*^+/-^;v*clb*^-/-^) embryos), **p<0.01, ***p<0.001 (One-way ANOVA and Dunnett’s post-test). **(H)** Violin plot showing average finger number per ISV of *vinculin* wild-type (*vcla*^+/+^;*vclb*^+/+^), single heterozygous (*vcla*^+/-^;*vclb*^+/+^), double heterozygous (*vcla*^+/-^;*vclb*^+/-^) or *vclb* homozygous knockout (*vcla*^+/-^ *;vclb^-/-^* embryos (n = 4 (*vcla*^+/+^;*vclb*^+/+^), 9 (*vcla*^+/-^;*vclb*^+/+^), 22 (*vcla*^+/-^;*vclb*^+/-^) and 12 *(vcla^+/-^;vclb^-/-^*) embryos), *** p<0.001 (Kruskal-Wallis Test and Dunnett’s post-test).

### Vinculin is required for ISV morphogenesis

To investigate if Vinculin plays a role in endothelial junctional finger formation, we took a genetic loss-of-function approach. The zebrafish genome contains two *vinculin* paralogs, *vinculin a (vcla)* and *vinculin b (vclb),* which share high sequence conservation with their mammalian orthologs including human *VCL* (87% and 86% identical amino acids, respectively) (Han et al., 2017). Zebrafish loss-of-function mutants for *vcla* and *vclb* have previously been published (Fukuda et al., 2019a) (Cheng et al., 2016; El-Brolosy et al., 2019; Han et al., 2017).

To investigate the consequence of Vinculin depletion for sprouting angiogenesis, we employed the *vcl* knockout models generated by Han et al. (2017) and crossed the *vcla^hu10818^* homozygous; *vclb^hu11202^* heterozygous *(vcla^-/-^;vlcb^+/-^)* fish in the Tg(*fli1a:EGFP^y1^*) vascular reporter line. Interestingly, live-imaging of the offspring revealed vascular defects in early angiogenic ISV sprouts of *vcl* double knockout *(vela^-/-^;vclb^-/-^*) and *vcl* heterozygous *(vcla^-/-^;vclb^+/-^* that appeared around the onset of lumen formation at 28 hpf (Supplementary Fig. 3). In wild-type *(vcla^+/+^;vclb^+/+^)* embryos, ISVs sprouted out of the dorsal aorta (DA) from 28 hpf stage onwards and once tip cells reached the dorsolateral roof of the neural tube, they anastomosed (Supplementary Fig. 3 A-A1). In contrast, in *vclb* heterozygous (*vcla^-/-^;vclb^+/-^*) and *vcl* double knockout (*vcla*^-/-^;*vclb*^-/-^) embryos, equal number of filopodial-forming tip cells and stalk cells were formed (Supplementary Fig. 3 A-C and Supplementary Fig. 4 A-E). Notably, we noticed a gradual increase in the number of wider ISV sprouts upon the deletion of *vinculin* (Supplementary Fig. 3 D and Supplementary. Fig. 4 A-E). This corresponded with a decreased ISV length and increased diameter in the *vcl* heterozygous and *vcl* double knockout embryos (Supplementary Fig. 3 E-F, Supplementary Fig. 4 F-G)), pointing towards changes in endothelial rearrangements upon *vinculin* deletion. This led to a delay of the angiogenesis process in the ISVs of the *vcl* double knockout embryos (Supplementary Fig. 3 and 4, Supplementary Movies 5-6). Moreover, ISVs were lumenized and there was blood flow in *vcl* double knockout, as was evident by the presence of erythrocytes within the developing blood vessel (Supplementary Fig. 5, Supplementary Movie 7).

### Endothelial Vinculin is required and sufficient for the formation of junctional fingers

To assess whether *vinculin* is required for the formation of junctional fingers, we performed live-imaging of *vcl* mutants expressing a junctional (Pecam1-EGFP) and a membrane (mCherry-CAAX) reporter (Fig. 5 D-F). In *vcl* double heterozygotes *(vcla^+/-^;vclb^+/-^* (Fig. 5 E), we observed a significant decrease in both the number and size of the junctional fingers upon lumen opening when compared to wild-type *(vcla^+/+^;vclb^+/+^) or* single heterozygous *(vcla^+/-^;vclb^+/+^*) (Fig. 5 D). Moreover, the homozygous loss of *vclb (vcla^+/-^;vclb^-/-^*) (Fig. 5 F) strongly reduced the formation of junctional fingers altogether. Interestingly, we observed gradual phenotypes of junctional finger size and length exerted by the various genotypes corresponding with the level of Vinculin deletion (Fig. 5 G-H).

Although the above findings demonstrate a strict requirement of Vinculin for junctional finger formation, these vascular defects may also be caused by potential secondary changes in other tissues in *vcl* mutants. We therefore wanted to test whether endothelial expression of vinculin is sufficient to restore junctional finger formation. To this end, we injected plasmid DNA encoding mCherry-Vinculin *(kdrl:mCherry-vinculin)* into *vcl* mutant embryos. This resulted in mosaic endothelial-specific expression of Vinculin in ISVs (Fig. 6 A-D). The expression of mCherry-Vinculin did not affect junctional finger formation in injected single heterozygous embryos *(vcla^+/-^;vclb^+/+^)* (Fig. 6 A-A’2), which phenotypically appear similar to the wild-type control group (Fig. 5 D, G-H). In contrast, and consistent with our earlier results, *vcl* double heterozygous *(ycla^+/-^;vclb^+/-^*) embryos displayed reduced junctional fingers in mCherry-Vinculin negative ISVs (Fig. 6 B1), whereas the ISVs expressing *kdrl:mCherry-vinculin* were able to form junctional fingers with their neighboring endothelial cells (Fig. 6 B2-B’2). Moreover, mCherry-Vinculin expression readily rescued junctional finger formation in homozygous *vclb* mutants *(vcla^+/-^;vclb^-/-^*), which otherwise did not form junctional fingers at all (Fig. 6 C-C’2). Taken together, these results clearly show that Vinculin is essential for the formation of junctional fingers and that expression of Vinculin within the endothelium is sufficient to promote finger formation in a *vcl*-knockout background.

**Figure 6.**
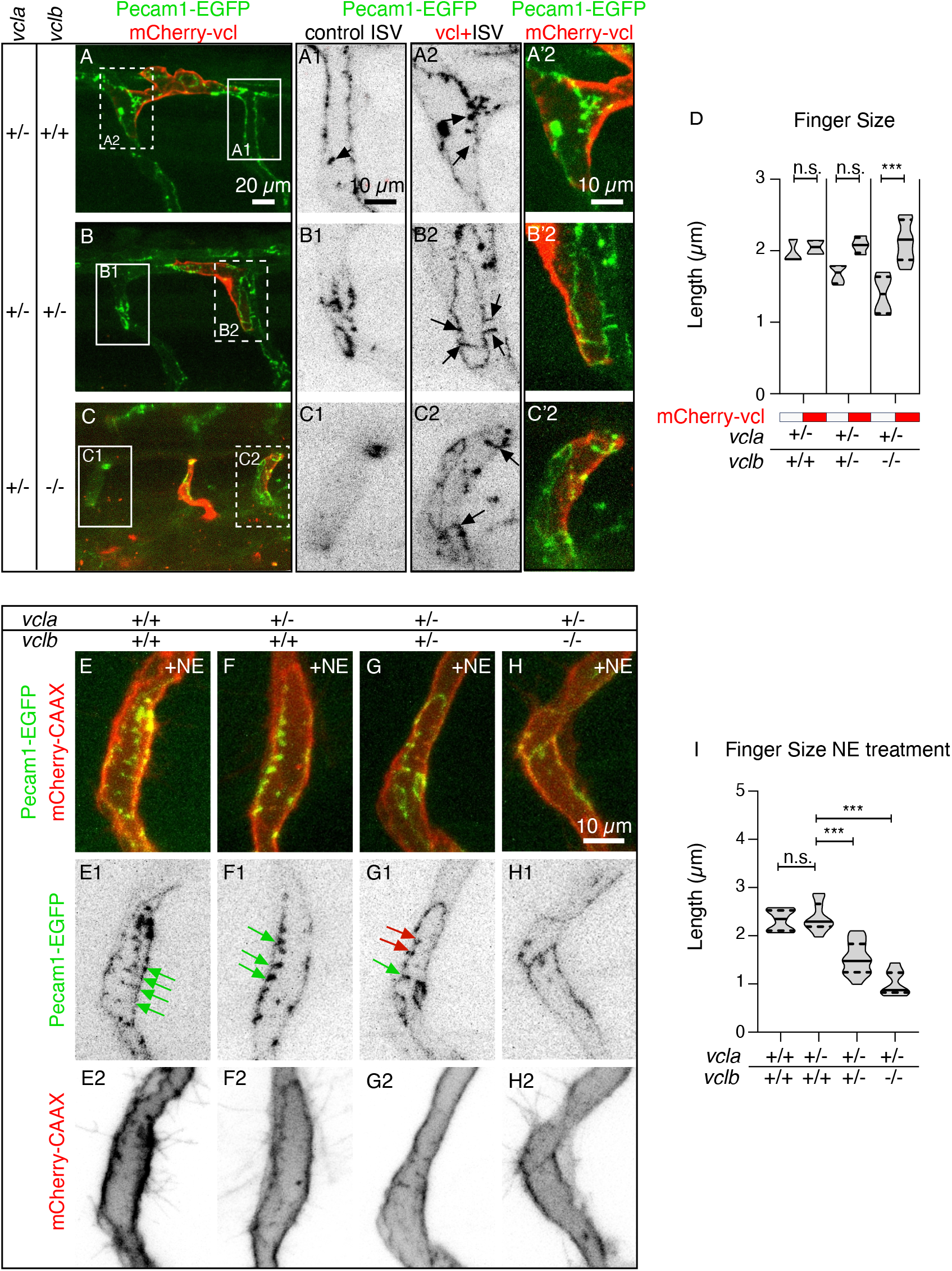
Increase in blood pressure does not rescue the loss of junctional fingers in *vinculin* mutants. **(A-C)** Confocal images of 36 hpf *Tg(fli:pecam1-EGFP)* embryos injected with *kdrl:mCherry-vinculin* with genotypes (A) single heterozygous (*vcla*^+/-^;*vclb*^+/+^), (B) double heterozygous mutant (*vcla*^+/-^;*vclb*^+/-^) or (C) *vclb* homozygous mutant (*vcla*^+/-^;*vclb*^-/-^). Scale bar, 20 μm. **(A1-C1)** Zoom-in inverted contrast images of *fli:pecam-EGFP* of the insets in AC of an ISV without *kdrl:mCherry-vinculin* expression (internal control). Scale bar, 10 μm. **(A2-C2)** Zoom-in inverted contrast images of *fli:pecam-EGFP* of the insets in A-C of an *flk:mCherry-vinculin* expressing ISV. **(A’2-C’2)** Merged image of A2-C2. Arrows indicate junctional fingers. **(D)** Violin plot showing the average junctional finger length per ISV ±S.E.M. of *vinculin* single heterozygous (*vcla*^+/-^;*vclb*^+/+^), double heterozygous mutant *(vcla^+/-^;vclb^+/-^* or *vclb* homozygous mutant *(vcla^+/-^;vclb^-/-^* embryos with or without kdrl:mCherry-*vinculin* expression. (n = 2 (*vcla*^+/-^;*vclb*^+/+^), 3 (*vcla*^+/-^;*vclb*^+/-^) and 5 (*vcla*^+/-^;*vclb*^-/-^) embryos), ***p<0.001 (Kruskal-Wallis Test and Dunnett’s post-test). **(E-H)** Confocal images of ISVs from *Tg(kdrl:mCherry-CAAX;fli:pecam1-EGFP)* embryos at 36 hpf with genotypes (E) wildtype (*vcla^+/+^;vclb*^+/+^), (F) single heterozygous (*vcla*^+/-^;*vclb*^+/+^); (G) double heterozygous mutant *(ycla^+/-^;vclb^+/-^* or (H) *vclb* homozygous mutant *(vcla^+/-^;vclb^-/-^* after treatment with 60 μm norepinephrine (NE). GFP+ signal marks Pecam-1 and RFP+ signal indicates the cell membrane. Scale bar, 10 μm. **(E1-H1)** Inverted contrast images of pecam1-EGFP of the outlined boxes in E. Green arrows indicate junctional fingers and red arrows mark shorter junctional fingers in the double heterozygous mutant *(vcla^+/-^;vclb^+/-^*. Note the absence of junctional fingers in *vclb* homozygous knock outs. **(E2-H2)** Inverted contrast images of mCherry-CAAX of the outlined boxes in E. **(I)** Violin plot showing the average junctional finger length per ISV after NE treatment of the genotypes wild-type (*vcla^+/+^;vclb*^+/+^), single heterozygous (*vcla*^+/-^;*vclb*^+/+^); double heterozygous mutant *(vcla^+/-^;vclb^+/-^* or *vclb* homozygous mutant *(vcla^+/-^;vclb^-/-^*). (n = 7 (*vcla^+/+^;vclb^+/+^*), 10 (*vcla*^+/-^;*vcla*^+/+^), 17 *(vcla^+/-^;vclb^+/-^* and 7 *(vcla^+/-^;vclb^-/-^* ISVs), n.s. = non-signficiant, *** p<0.001 (One-Way ANOVA with Dunnett’s post-test).

### Increase in blood pressure does not rescue the loss of junctional fingers in vinculin double knockouts

As shown above, the formation of junctional fingers is triggered by an increase in blood pressure. We therefore hypothesized that loss of *vinculin* may reduce the junctional response to hemodynamic forces and that this reduced sensitivity may be overcome by an increase in blood pressure. To test this possibility, we treated *vcl* double knockout embryos with NE to increase blood pressure at the onset of ISV lumen inflation. Notably, NE treatments did not increase the formation of junctional fingers in *vcl* heterozygous or homozygous mutants, in contrast to single heterozygous embryos, in which NE was able to enhance junctional fingers (Fig. 6 E-I). Thus, in *vcl* double knockouts, endothelial cells appear unresponsive to increased blood pressure. Furthermore, the effect was most pronounced in double mutants whereas combinations of mutant and wild-type alleles displayed an intermediate phenotype, suggesting a dose-dependent requirement of Vinculin in this process. Taken together, these results point to a mechanism in which Vinculin recruitment acts as an adaptive response to elevated junctional tension during angiogenic sprouting.

## Discussion

Dynamic junctional remodeling is essential for blood vessel formation, maintenance and function. Here, we describe the formation of junctional fingers as a novel trait of junctional remodeling which occurs during lumen formation of developing ISVs in the zebrafish embryo. Junctional fingers appear as junctional folds of up to 3-4 μm, that are orientated perpendicular to the originating junction and show a characteristic dynamic behavior. They are formed upon the interconnection between the dorsal aorta and the sprouting ISV, which triggers exposure of sprouting endothelial cells to blood pressure. However, junctional fingers disappear once ISVs have achieved patency and blood flow is firmly established. This specific response to blood pressure renders junctional fingers as a novel *in vivo* paradigm for the study of junctional dynamics in response to extrinsic forces.

Our studies demonstrate that formation of junctional fingers relies on the recruitment of Vinculin and thus establish an interconnection between junctional tension, junctional remodeling and blood vessel morphogenesis. Recent studies have established distinct roles for *vcla* and *vclb* during zebrafish heart development, which reflect their expression in the myocardium and epicardium, respectively (Cheng et al., 2016; Fukuda et al., 2019b; Han et al., 2017). In contrast, in the endothelium *vcla* and *vclb* are co-expressed (Lawson et al., 2020) and, consistent with this, we did not notice phenotypic variation in different mutant allele combinations (e.g. *vcla^-/-^;vclab^+/-^* vs. *vcla^+/-^;vclab^-/-^)* with regard to the appearance of junctional fingers - in agreement with the notion that *vcla* and *vclb* mutants can genetically compensate for each other (El-Brolosy et al., 2018).

Dynamic changes in junctional organization have been studied previously in different contexts. In particular, perpendicular-oriented endothelial cell-cell junctions have been described as serrated junctions, discontinuous junctions, focal adherens junctions (FAJs) and VE-cadherin fingers (Bentley et al., 2014; Hayer et al., 2016; Huveneers et al., 2012). Serrated and discontinuous junctions have been associated with active or remodeling junctions (Bentley et al., 2014). The junctional fingers we describe here appear as invaginations of stable and continuous junctions, and are morphologically similar to VE-cadherin fingers and FAJs that have been previously described in HUVECs (Huveneers et al., 2012) (Dorland and Huveneers, 2016; Hayer et al., 2016). However, because of the presence of Pecam1 and other cell-cell junctional components (ZO1, VE-cadherin, F-actin), we call them “junctional fingers” rather than “VE-cadherin fingers” in the current study.

VE-cadherin fingers and FAJs are involved in mechanical coupling between endothelial cells and are controlled by actomyosin contractility (Huveneers et al., 2012) (Dorland and Huveneers, 2016; Hayer et al., 2016). In agreement with this, junctional fingers form when sprouting endothelial cells encounter blood pressure. Modulation of blood flow by pharmacological intervention leads to a graded response, with more and longer junctional fingers correlating with higher blood pressure, suggesting that junctional fingers are an adaptive response of endothelial cells to increased mechanical stress. In fact, the specific localization of Vinculin to junctional fingers indicates that they are experiencing elevated junctional tension on the VE-cadherin-catenin complex (Huveneers et al., 2012). Our genetic analyses show that Vinculin is required for the formation of junctional fingers and mosaic expression in sprouting endothelial cells can rescue defects arising from *vinculin* loss-of function or *vinculin* knockout. Our previous cell culture experiments have shown that tension-induced Vinculin localization to FAJs protects these junctions against mechanical stress (Huveneers et al., 2012). Our *in vivo* observations suggest that Vinculin may have a dual role during vascular lumen inflation. First, a local increase in junctional tension triggers recruitment of Vinculin to the endothelial junction, which subsequently leads to the formation of junctional fingers. Alternatively, Vinculin may become localized upon junctional finger formation. In agreement with our study, recruitment of Vinculin-a to myocardial cell-cell junctions depends on the mechanical force generated by myocardial contractility (Fukuda et al., 2019b). Here, activated Vinculin recruits actin regulators and promotes myofiber maturation. It will be of interest to discern the molecular pathways that regulate Vinculin recruitment and function during vascular lumen expansion.

Junctional fingers show a dynamic behavior. They form during lumen expansion in sprouting vessels, but regress once blood flow is established. A likely scenario is that lumen inflation increases junctional tension, which is subsequently relieved by the commencement of blood flow. Further studies will be required to discern whether shear stress impacts on junctional finger maintenance or regression. However, we did not detect junctional fingers in the dorsal aorta or the posterior cardinal vein. These blood vessels are formed by vasculogenesis, which is likely to preclude junctional finger formation because of different hemodynamic conditions.

During vascular morphogenesis as well as in functional blood vessels, endothelial cells generate dynamic junctional protrusions, called junction-based lamellipodia (JBL) or junction-associated intermittent lamellipodia (JAIL), which have been observed in different vascular beds of zebrafish embryo as well as mouse embryos and the postnatal retina (reviewed by Cao and Schnittler, 2019; Fonseca et al., 2020). Live-imaging of JBL in zebrafish embryos showed that they are highly dynamic oscillatory structures and are associated with cell movements within the endothelium (Paatero et al., 2018; reviewed by Okuda and Hogan, 2020). Thus, JBL form in regions of high junctional turnover and reduced junctional stability. In contrast, the junctional fingers we describe here are transient, but do not oscillate. Their dynamic behavior and association with Vinculin indicate that they are strengthened junctions, which support junctional stability to maintain blood vessel integrity in response to sudden changes in extrinsic forces.

In conclusion, our results support a model (see Figure 7), in which endothelial cell-cell junctions form finger-like invaginations during vascular lumen expansion in response to junctional stretch induced by an increase of blood pressure. Junctional fingers recruit Vinculin and rely on it for their formation and stabilization. Taken together, junctional fingers represent a novel paradigm to study endothelial cell dynamics and junctional remodeling in response to mechanical forces during blood vessel morphogenesis *in vivo.*

**Figure 7.**
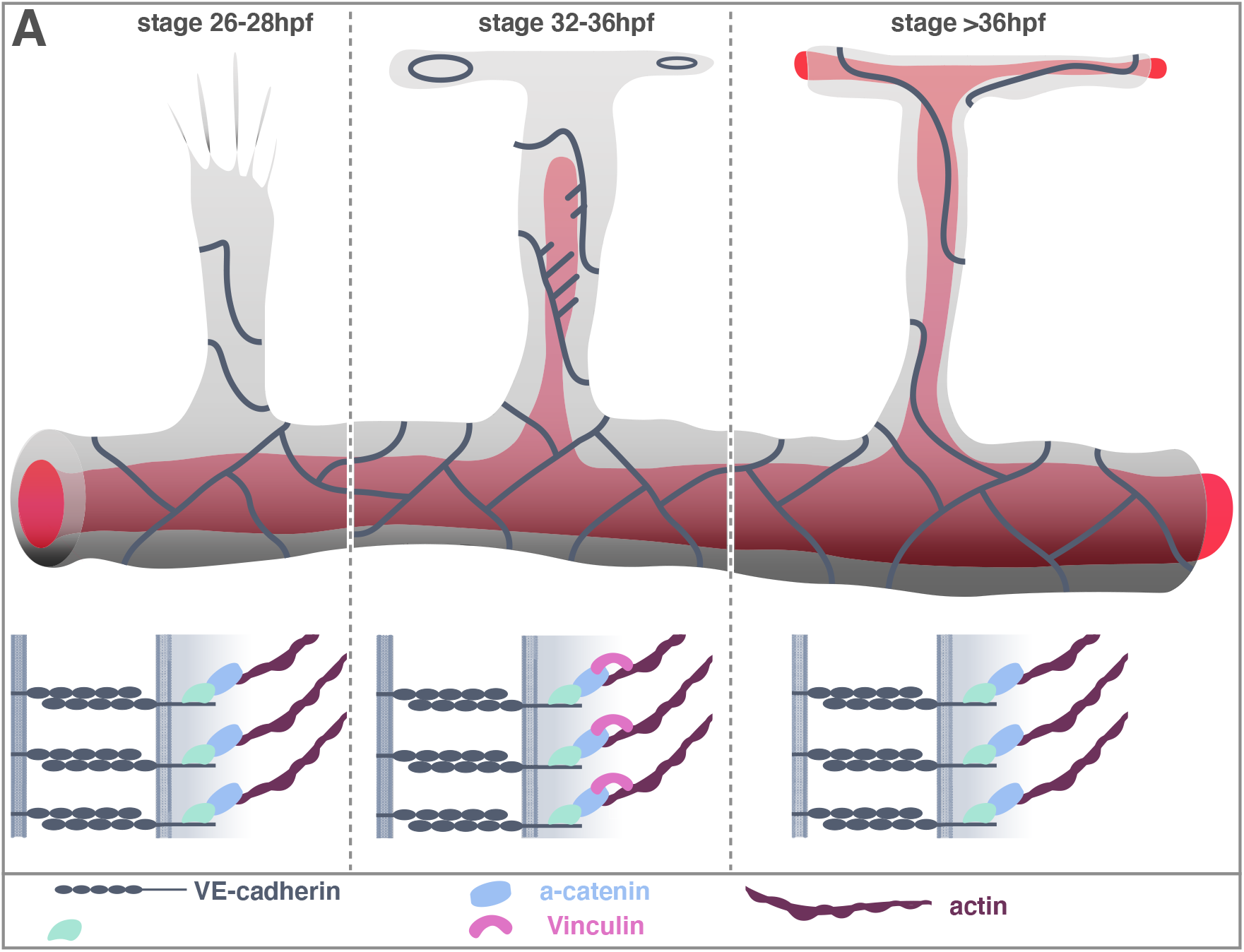
Summary Model. (A) During ISV sprouting (26-28hpf) endothelial cell junctions appear straight. Cell rearrangement lead to a multicellular configuration (B) Lumen expansion (32-36hpf). Once the lumen is opened endothelial cells in the nascent sprout become exposed to blood pressure. Junctional fingers occur in regions facing the expanding lumen. Vinculin is localized at junctional fingers. At this stage the ISV not yet patent. (C) Commencement of blood flow (40 hpf). Anastomosis generates a continuous vascular lumen permitting blood flow. Junctional fingers disappear, junctional Vinculin is not detectable.

## Materials and Methods

### Experimental Model and Subject Details

#### Zebrafish lines and maintenance

Zebrafish were maintained in standard conditions (Aleström et al., 2019). All experiments were performed in accordance with federal guidance and approved by the Kantonales Veterinäramt of Kanton Basel-Stadt. The vcla^*hu10818*^;vclb^*hu11202*^ (Han et al., 2017) zebrafish lines were crossed into the following transgenic lines: *Tgfli1a:EGFP)^□1^* (Lawson and Weinstein, 2002); *Tg(gata1a:DsRed)^sd2^* (Traver et al., 2003); *Tg(fli1a:pecam1-EGFP)^ncv27^* (Ando et al., 2016); *Tg(fli1a:GFF)^ubs3^* and *Tg(UAS:EGFP-hZO1)^ubs5^* (Herwig et al., 2011); *(Tg(UAS:mRuby2-UCHD)^ubs20^* (Paatero et al., 2018); *Tg(kdrl:EGFP-nls)^ubs1^* (Blum et al., 2008); *Tg(kdrl:mCherry-CAAX)^s916^* (Hogan et al., 2009); *Tg(fli1a:Lifeact-EGFP)* (Phng et al., 2013). Experiments were performed between 24 hours post fertilization (hpf) and 48 hpf. Before experiments, embryos were selected for fluorescent signal. After the experiments with the vinculin mutants, genotype was assessed by DNA extraction and PCR.

### Method Details

#### Genotyping of the vcl mutant lines

Genomic DNA from fin biopsies from adult fish or whole embryos was extracted by incubating the biopsies with H_2_O with addition of proteinase K for 30 minutes at 37°C. Subsequently, NaOH was added to a final concentration of 50 mM and incubated for 10 minutes at 95°C. Finally, 1/10^th^ of 1M Tris HCl pH 8 was added (Meeker et al., 2007). Genomic DNA was then used to genotype the *vcla* and *vclb* loci. For genotyping analysis, four different primers were used to distinguish between wild-type, heterozygous or homozygous *vcla* or *vclb* alleles. **For *vcla***, combination of four primers (FP 5’-AGCTGATGATCTGAATCAGGTGTG-3’, RP 5’-TCGTTCAATCACTCGTGCAAACAG-3’, Fwd-wt 5’-ACCATGGAGGACTTGATCACT-3’, Rev-mut 5’-GGTCCCAGGTTTTTAGTGTAAGT-3’) generated PCR products of 497 bp and 203 bp for the wild-type, three PCR products of 497 bp, 282 bp and 203 bp for the heterozygous *vcla* mutants and for PCR products of 497 bp and 282 bp for homozygous vcla mutants. **For *vclb***, combination of four primers (FP 5’-CGGTAGTTAGTTAGTTGTAGAGGGAGTC-3’, RP 5’-AATGAGAAAGCCTGAATGTGCG-3’, Rev-wt 5’-GCATGATCACCAGATGGG-3’, Fwd-mut 5’-CCCAGCAGATCTGGTGAT-3’) generated PCR products of 490 bp and 201 bp for the wild-type, PCR products of 490 bp, 319 bp and 201 for the heterozygous *vclb* mutants and PCR products of 490 bp and 319 bp for homozygous *vclb* mutants PCR was performed using OneTaq DNA polymerase (NEB) and PCR products were loaded on a 1% agarose gel to assess the genotype (see Supplementary Figure 2).

#### Generation of kdrl:mCherry-vinculin expression plasmid

In a first step the kdrl promoter (Genbank accession number: AY045466) was subcloned in the tol2 transposon plasmid pDB739 (Balciunas et al., 2006) using the restriction sites BamHI and EcoRV. The N-terminal tagged mCherry-vinculin open reading frame (Twiss et al., 2012) was then inserted into the EcoRV restriction site adjacent to the kdrl promoter.

#### Embryo Microinjections

For transient expression of the kdrl:mCherry-vinculin, plasmid DNA was injected using glass needles (Biomedical instruments) and standard microinjection protocol (Lenard et al., 2013) in (*Tgfli1a:pecam1-EGFP)^ncv27^* or *vcla^hu10818^;vclb^hu1202^Tgfli1a:pecam1-EGFP)^ncv27^* embryos at the one-or two-cell stage.

#### Immunofluorescence

Immunofluorescence staining of embryos was performed as previously described (Herwig et al., 2011) with few modifications. Briefly, embryos were fixed in 2% PFA in PBST (0.2% tween in PBS) and incubated overnight at 4°C. Embryos were washed in PBST and consecutively permeabilized in 0,5% Triton/PBST for 30-45 minutes at room temperature, and subsequently blocked (1% BSA, 0.2% Triton, 5% Goat Serum, 0,01% sodium azide in PBST) overnight at 4°C. Embryos were stained with mouse-anti-human-ZO-1 1:500 (Thermo Fisher Scientific, 33-9100) or guinea pig-anti-zebrafish-VE-cadherin 1:500 (Paatero, 2018) diluted in Pierce Immunostain Enhancer medium (Thermo Fisher Scientific). Alexa Fluor 405 goat anti-mouse immunoglobulin (IgG) 1:500 (Thermo Fisher Scientific) and Alexa Fluor 568 goat anti-guinea pig IgG 1:500 (Thermo Fisher Scientific) were used as secondary antibodies.

#### Imaging and Image Analysis

Fixed or live embryos were selected for fluorescence signal, anaesthetized in E3 fish water (5 mM NaCl, 0.17 mM KCl, 0.33 mM CaCl2, 0.33 mM MgSO4, pH 7.4) with 1x tricaine (0.08%, Sigma) and mounted in glass bottom Petri dishes (MatTek) using 0.7% low-melting-point agarose (Sigma) containing 1x tricaine. For live-imaging, E3 with 1x tricaine and 0.003% 1-phenyl-2-thiourea (PTU, Sigma) to avoid pigmentation, was added to the dish. A Zeiss LSM880 Airyscan inverted confocal microscope was used for live-imaging and imaging of fixed samples. For fixed samples, the 40x (NA = 1.3) silicon oil objective was used. Z-stacks were made with a step size of 0.5 μm. For live imaging, the 25x (NA = 0.8) oil objective was used. Images were acquired with a zoom of 1-1.6 and z-stacks were made with a step size of 0.5 to 1.0 μm. Frames were acquired every 5-30 minutes. High resolution timelapse images were acquired with an Olympus SpinSR spinning disc microscope using a 30x (NA = 1.05) oil objective (Photometrics). Frames were acquired every 1-2 minutes with a z-stack step size of 0.7 μm. Images were analyzed with the ImageJ software.

#### Pharmacological Treatments

To reduce heartbeat and inhibit blood circulation, we treated zebrafish embryos with 4x tricaine (0.32%, Sigma) in E3 embryo medium with 0.003% PTU (Sigma), as previously described (Lenard et al., 2013). To increase heartbeat and blood flow, we treated zebrafish embryos with 60 μM norepinephrine (Sigma) in E3 fish water containing 0.003% PTU, as previously described (Chen et al., 2012; De Luca et al., 2014). Treatment of tricaine or norepinephrine started directly during imaging as indicated in the corresponding figures at 30-32 hpf.

#### Quantification and Statistical Analysis

Images were analyzed with the ImageJ software. For analysis of blood vessel length, an average of three measurements was taken for each ISV and for blood vessel diameter, an average of three measurements was taken at different levels of each ISV. For analysis of the finger number, averages were taken for each ISV and junctional finger size was traced in three measurements per finger and averaged per ISV. Microsoft Excel was used for data analysis and Prism Graphpad V6 was used for statistical analysis and data visualization. Violin plots represent median ±quartiles. When 2 groups were compared, a Student’s T-test was used. A One-way analysis of variance (ANOVA) was used when 2 or more groups were compared to the control, in combination with a Dunnett’s test for multiple comparisons and a D’Agostino-Pearson test for normality. Asterisks indicate p values, and are defined as n.s. Non-significant, * p<0.05, ** p <0.01, *** p<0.001.

## Acknowledgements

We thank Kumuthini Kulendra for fish care and the Imaging Core Facility of the Biozentrum (University of Basel) for microscopy support. This work has been supported by the Kantons Basel-Stadt and Basel-Land and by a grant from the Swiss National Science Foundation to M.A.. S.H. is financially supported by the Netherlands Organization of Scientific Research (NWO)-ZonMw VIDI grant 016.156.327). M.S. was financially supported by (EUFish, UvA365, Amsterdam UMC).

## Author Contributions

H.G.B. and S.H. conceived the idea and directed the work. M.A. provided supervision and financial support. M.P.K. and M.S. designed and performed experiments. H.G.B., S.H., M.A., M.P.K. and M.S. analyzed the data. M.K.H., B.K., and J.d.R. provided essential reagents. M.P.K. and M.S. prepared figures. H.G.B., S.H., M.P.K. and M.S. wrote the manuscript. All authors reviewed the manuscript.

## References

Aleström, P., D’Angelo, L., Midtlyng, P.J., Schorderet, D.F., Schulte-Merker, S., Sohm, F., Warner, S., 2019. Zebrafish: Housing and husbandry recommendations. Lab Anim 51, 002367721986903–12. doi:10.1177/0023677219869037

Ando, K., Fukuhara, S., Izumi, N., Nakajima, H., Fukui, H., Kelsh, R.N., Mochizuki, N., 2016. Clarification of mural cell coverage of vascular endothelial cells by live imaging of zebrafish. Development 143, 1328–1339. doi:10.1242/dev.132654

Angulo-Urarte, A., van der Wal, T., Huveneers, S., 2020. Cell-cell junctions as sensors and transducers of mechanical forces. BBA - Biomembranes 1862, 183316. doi:10.1016/j.bbamem.2020.183316

Baeyens, N., Schwartz, M.A., 2016. Biomechanics of vascular mechanosensation and remodeling. MBoC 27, 7–11. doi:10.1091/mbc.E14-11-1522

Balciunas, D., Wangensteen, K.J., Wilber, A., Bell, J., Geurts, A., Sivasubbu, S., Wang, X., Hackett, P.B., Largaespada, D.A., McIvor, R.S., Ekker, S.C., 2006. Harnessing a high cargo-capacity transposon for genetic applications in vertebrates. PLoS Genet 2, e169. doi:10.1371/journal.pgen.0020169

Barry, A.K., Wang, N., Leckband, D.E., 2015. Local VE-cadherin mechanotransduction triggers long-ranged remodeling of endothelial monolayers. Journal of Cell Science 128, 1341–1351. doi:10.1242/jcs.159954

Bentley, K., Franco, C.A., Philippides, A., Blanco, R., Dierkes, M., Gebala, V., Stanchi, F., Jones, M., Aspalter, I.M., Cagna, G., Weström, S., Claesson-Welsh, L., Vestweber, D., Gerhardt, H., 2014. The role of differential VE-cadherin dynamics in cell rearrangement during angiogenesis. Nat Cell Biol 16, 309–321. doi:10.1038/ncb2926

Betz, C., Lenard, A., Belting, H.-G., Affolter, M., 2016. Cell behaviors and dynamics during angiogenesis. Development 143, 2249–2260. doi:10.1242/dev.135616

Blum, Y., Belting, H.-G., Ellertsdottir, E., Herwig, L., Lüders, F., Affolter, M., 2008. Complex cell rearrangements during intersegmental vessel sprouting and vessel fusion in the zebrafish embryo. Developmental Biology 316, 312–322. doi:10.1016/j.ydbio.2008.01.038

Cao, J., Schnittler, H., 2019. Putting VE-cadherin into JAIL for junction remodeling. Journal of Cell Science 132, jcs222893–19. doi:10.1242/jcs.222893

Chen, Q., Jiang, L., Li, C., Hu, D., Bu, J.-W., Cai, D., Du, J.-L., 2012. Haemodynamics-Driven Developmental Pruning of Brain Vasculature in Zebrafish. PLoS Biol 10, e1001374–18. doi:10.1371/journal.pbio.1001374

Cheng, F., Miao, L., Wu, Q., Gong, X., Xiong, J., Zhang, J., 2016. Vinculin b deficiency causes epicardial hyperplasia and coronary vessel disorganization in zebrafish. Development 143, 3522–3531. doi:10.1242/dev.132936

Conway, D.E., Breckenridge, M.T., Hinde, E., Gratton, E., Chen, C.S., Schwartz, M.A., 2013. Fluid Shear Stress on Endothelial Cells Modulates Mechanical Tension across VE-Cadherin and PECAM-1. Current Biology 23, 1024–1030. doi:10.1016/j.cub.2013.04.049

De Luca, E., Zaccaria, G.M., Hadhoud, M., Rizzo, G., Ponzini, R., Morbiducci, U., Santoro, M.M., 2014. ZebraBeat: a flexible platform for the analysis of the cardiac rate in zebrafish embryos. Scientific Reports 4, 835–13. doi:10.1038/srep04898

Dorland, Y.L., Huveneers, S., 2016. Cell-cell junctional mechanotransduction in endothelial remodeling. Cellular and Molecular Life Sciences 1–14. doi:10.1007/s00018-016-2325-8

Duchemin, A.-L., Vignes, H., Vermot, J., Chow, R., 2019. Mechanotransduction in cardiovascular morphogenesis and tissue engineering. Current Opinion in Genetics & Development 57, 106–116. doi:10.1016/j.gde.2019.08.002

El-Brolosy, M.A., Kontarakis, Z., Rossi, A., Kuenne, C., nther, S.G.X., Fukuda, N., Kikhi, K., Boezio, G.L.M., Takacs, C.M., Lai, S.-L., Fukuda, R., Gerri, C., Giraldez, A.J., Stainier, D.Y.R., 2019. Genetic compensation triggered by mutant mRNA degradation. Nature Publishing Group 1–26. doi:10.1038/s41586-019-1064-z

El-Brolosy, M.A., Rossi, A., Kontarakis, Z., Kuenne, C., Günther, S., Fukuda, N., Takacs, C., Lai, S.-L., Fukuda, R., Gerri, C., Kikhi, K., Giraldez, A.J., Stainier, D.Y.R., 2018. Genetic compensation is triggered by mutant mRNA degradation 83, 2031–29. doi:10.1101/328153

Fonseca, C.G., Barbacena, P., Franco, C.A., 2020. Endothelial cells on the move: dynamics in vascular morphogenesis and disease. Vascular Biology 2, H29–H43. doi:10.1530/VB-20-0007

Fukuda, R., Gunawan, F., Ramadass, R., Beisaw, A., Konzer, A., Mullapudi, S.T., Gentile, A., Maischein, H.-M., Graumann, J., Stainier, D.Y.R., 2019a. Mechanical Forces Regulate Cardiomyocyte Myofilament Maturation via the VCL-SSH1-CFL Axis. Developmental Cell 51, 62–77.e5. doi:10.1016/j.devcel.2019.08.006

Fukuda, R., Gunawan, F., Ramadass, R., Beisaw, A., Konzer, A., Mullapudi, S.T., Gentile, A., Maischein, H.-M., Graumann, J., Stainier, D.Y.R., 2019b. Mechanical Forces Regulate Cardiomyocyte Myofilament Maturation via the VCL-SSH1-CFL Axis. DEVCEL 51, 62–77.e5. doi:10.1016/j.devcel.2019.08.006

Gebala, V., Collins, R., Geudens, I., Phng, L.-K., Gerhardt, H., 2016. Blood flow drives lumen formation by inverse membrane blebbing during angiogenesis in vivo. Nat Cell Biol 18, 443–450. doi:10.1038/ncb3320

Han, M.K.L., van der Krogt, G.N.M., de Rooij, J., 2017. Zygotic vinculin is not essential for embryonic development in zebrafish. PLoS ONE 12, e0182278. doi:10.1371/journal.pone.0182278

Hayer, A., Shao, L., Chung, M., Joubert, L.-M., Yang, H.W., Tsai, F.-C., Bisaria, A., Betzig, E., Meyer, T., 2016. Engulfed cadherin fingers are polarized junctional structures between collectively migrating endothelial cells. Nat Cell Biol 18, 1311–1323. doi:10.1038/ncb3438

Herwig, L., Blum, Y., Krudewig, A., Ellertsdottir, E., Lenard, A., Belting, H.-G., Affolter, M., 2011. Distinct Cellular Mechanisms of Blood Vessel Fusion in the Zebrafish Embryo. Current Biology 21, 1942–1948. doi:10.1016/j.cub.2011.10.016

Hoefer, I.E., Adel, den, B., Daemen, M.J.A.P., 2013. Biomechanical factors as triggers of vascular growth. Cardiovascular Research 99, 276–283. doi:10.1093/cvr/cvt089

Hogan, B.M., Bos, F.L., Bussmann, J., Witte, M., Chi, N.C., Duckers, H.J., Schulte-Merker, S., 2009. ccbe1 is required for embryonic lymphangiogenesis and venous sprouting. Nat Genet 41, 396–398. doi:10.1038/ng.321

Hogan, B.M., Schulte-Merker, S., 2017. How to Plumb a Pisces: Understanding Vascular Development and Disease Using Zebrafish Embryos. DEVCEL 42, 567–583. doi:10.1016/j.devcel.2017.08.015

Huveneers, S., Oldenburg, J., Spanjaard, E., van der Krogt, G., Grigoriev, I., Akhmanova, A., Rehmann, H., de Rooij, J., 2012. Vinculin associates with endothelial VE-cadherin junctions to control force-dependent remodeling. The Journal of Cell Biology 196, 641–652. doi:10.1083/jcb.201108120

Komarova, Y.A., Kruse, K., Mehta, D., Malik, A.B., 2017. Protein Interactions at Endothelial Junctions and Signaling Mechanisms Regulating Endothelial Permeability. Circ Res 120, 179–206. doi:10.1161/CIRCRESAHA.116.306534

Lagendijk, A.K., Gomez, G.A., Baek, S., Hesselson, D., Hughes, W.E., Paterson, S., Conway, D.E., Belting, H.-G., Affolter, M., Smith, K.A., Schwartz, M.A., Yap, A.S., Hogan, B.M., 2017. Live imaging molecular changes in junctional tension upon VE-cadherin in zebrafish. Nature Communications 1–12. doi:10.1038/s41467-017-01325-6

Lampugnani, M.G., Dejana, E., Giampietro, C., 2017. Vascular Endothelial (VE)-Cadherin, Endothelial Adherens Junctions, and Vascular Disease (2017). Cold Spring Harbor Perspectives in Biology 9. doi:10.1101/cshperspect.a033720

Lawson, N.D., Li, R., Shin, M., Grosse, A., Yukselen, O., Stone, O.A., Kucukural, A., Zhu, L., 2020. An improved zebrafish transcriptome annotation for sensitive and comprehensive detection of cell type-specific genes. eLife 9, 347–28. doi:10.7554/eLife.55792

Lawson, N.D., Weinstein, B.M., 2002. In Vivo Imaging of Embryonic Vascular Development Using Transgenic Zebrafish. Developmental Biology 248, 307–318. doi:10.1006/dbio.2002.0711

Lenard, A., Ellertsdottir, E., Herwig, L., Krudewig, A., Sauteur, L., Belting, H.-G., Affolter, M., 2013. In vivo analysis reveals a highly stereotypic morphogenetic pathway of vascular anastomosis. Developmental Cell 25, 492–506. doi:10.1016/j.devcel.2013.05.010

Liu, Z., Tan, J.L., Cohen, D.M., Yang, M.T., Sniadecki, N.J., Ruiz, S.A., Nelson, C.M., Chen, C.S., 2010. Mechanical tugging force regulates the size of cell-cell junctions. Proc. Natl. Acad. Sci. U.S.A. 107, 9944–9949. doi:10.1073/pnas.0914547107

Meeker, N.D., Hutchinson, S.A., Ho, L., Trede, N.S., 2007. Method for isolation of PCR-ready genomic DNA from zebrafish tissues. Biotechniques 43, 610–612-614. doi:10.2144/000112619

Okuda, K.S., Hogan, B.M., 2020. Endothelial Cell Dynamics in Vascular Development: Insights From Live-Imaging in Zebrafish. Front Physiol 11, 842. doi:10.3389/fphys.2020.00842

Paatero, I., Sauteur, L., Lee, M., Lagendijk, A.K., Heutschi, D., Wiesner, C., Guzmán, C., Bieli, D., Hogan, B.M., Affolter, M., Belting, H.-G., 2018. Junction-based lamellipodia drive endothelial cell rearrangements in vivo via a VE-cadherin-F-actin based oscillatory cell-cell interaction. Nature Communications 1–13. doi:10.1038/s41467-018-05851-9

Phng, L.-K., Stanchi, F., Gerhardt, H., 2013. Filopodia are dispensable for endothelial tip cell guidance. Development 140, 4031–4040. doi:10.1242/dev.097352

Potente, M., Gerhardt, H., Carmeliet, P., 2011. Basic and Therapeutic Aspects of Angiogenesis. Cell 146, 873–887. doi:10.1016/j.cell.2011.08.039

Roman, B.L., Pekkan, K., 2012. Mechanotransduction in embryonic vascular development. Biomech Model Mechanobiol 11, 1149–1168. doi:10.1007/s10237-012-0412-9

Sauteur, L., Krudewig, A., Herwig, L., Ehrenfeuchter, N., Lenard, A., Affolter, M., Belting, H.-G., 2014. Cdh5/VE-cadherin Promotes Endothelial Cell Interface Elongation via Cortical Actin Polymerization during Angiogenic Sprouting. CellReports 9, 504–513. doi:10.1016/j.celrep.2014.09.024

Szymborska, A., Gerhardt, H., 2018. Hold Me, but Not Too Tight—Endothelial Cell-Cell Junctions in Angiogenesis. Cold Spring Harbor Perspectives in Biology 10, a029223–17. doi:10.1101/cshperspect.a029223

Traver, D., Paw, B.H., Poss, K.D., Penberthy, W.T., Lin, S., Zon, L.I., 2003. Transplantation and in vivo imaging of multilineage engraftment in zebrafish bloodless mutants. Nat Immunol 4, 1238–1246. doi:10.1038/ni1007

Twiss, F., le Duc, Q., Van Der Horst, S., Tabdili, H., van der Krogt, G., Wang, N., Rehmann, H., Huveneers, S., Leckband, D.E., de Rooij, J., 2012. Vinculin-dependent Cadherin mechanosensing regulates efficient epithelial barrier formation. Biology Open 1, 1128–1140. doi:10.1242/bio.20122428

Tzima, E., Irani-Tehrani, M., Kiosses, W.B., Dejana, E., Schultz, D.A., Engelhardt, B., Cao, G., DeLisser, H., Schwartz, M.A., 2005. A mechanosensory complex that mediates the endothelial cell response to fluid shear stress. Nature 437, 426–431. doi:10.1038/nature03952

Wallez, Y., Huber, P., 2008. Endothelial adherens and tight junctions in vascular homeostasis, inflammation and angiogenesis. Biochimica et Biophysica Acta (BBA) - Biomembranes 1778, 794–809. doi:10.1016/j.bbamem.2007.09.003

Yao, M., Qiu, W., Liu, R., Efremov, A.K., Cong, P., Seddiki, R., Payre, M., Lim, C.T., Ladoux, B., ge, R.E.M.M.E., Yan, J., 2014. Force-dependent conformational switch of alpha-catenin controls vinculin binding. Nature Communications 1–12. doi:10.1038/ncomms5525

