## Supplementary figures and images for "Vinculin controls endothelial cell junction dynamics during vascular lumen formation"

### Supplementary Figures 1-5

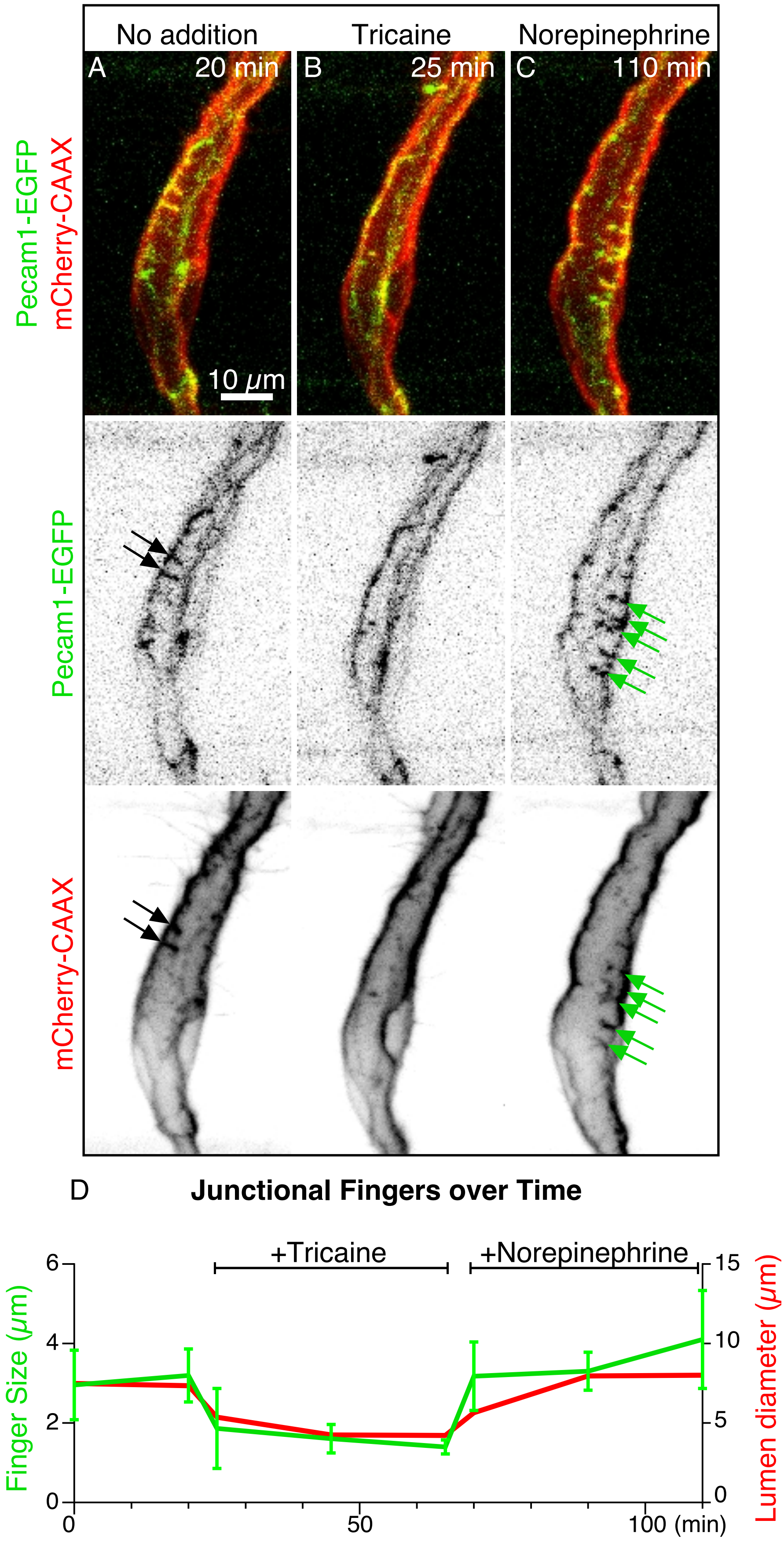


#### Suppl.Figure1

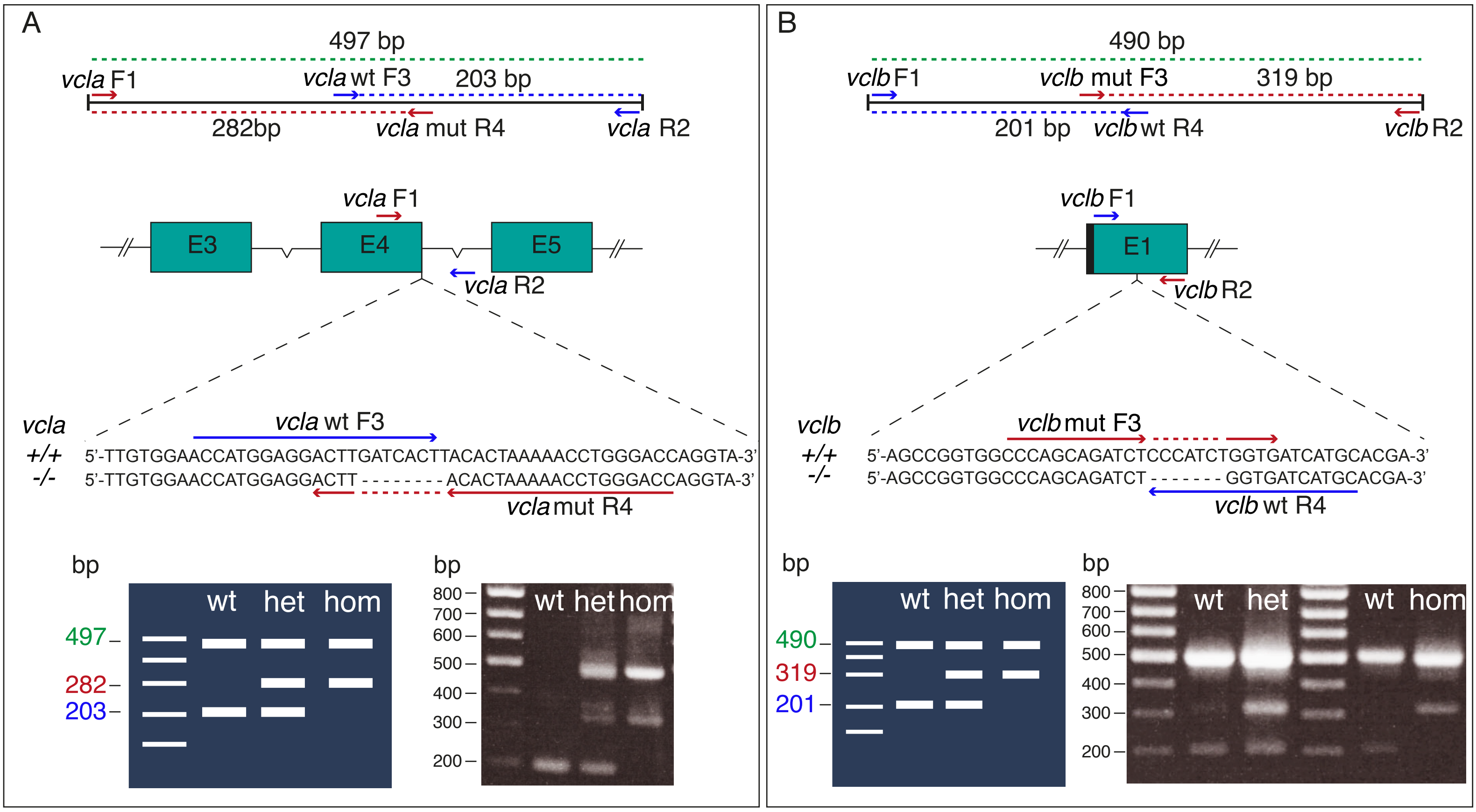


#### Suppl.Figure2

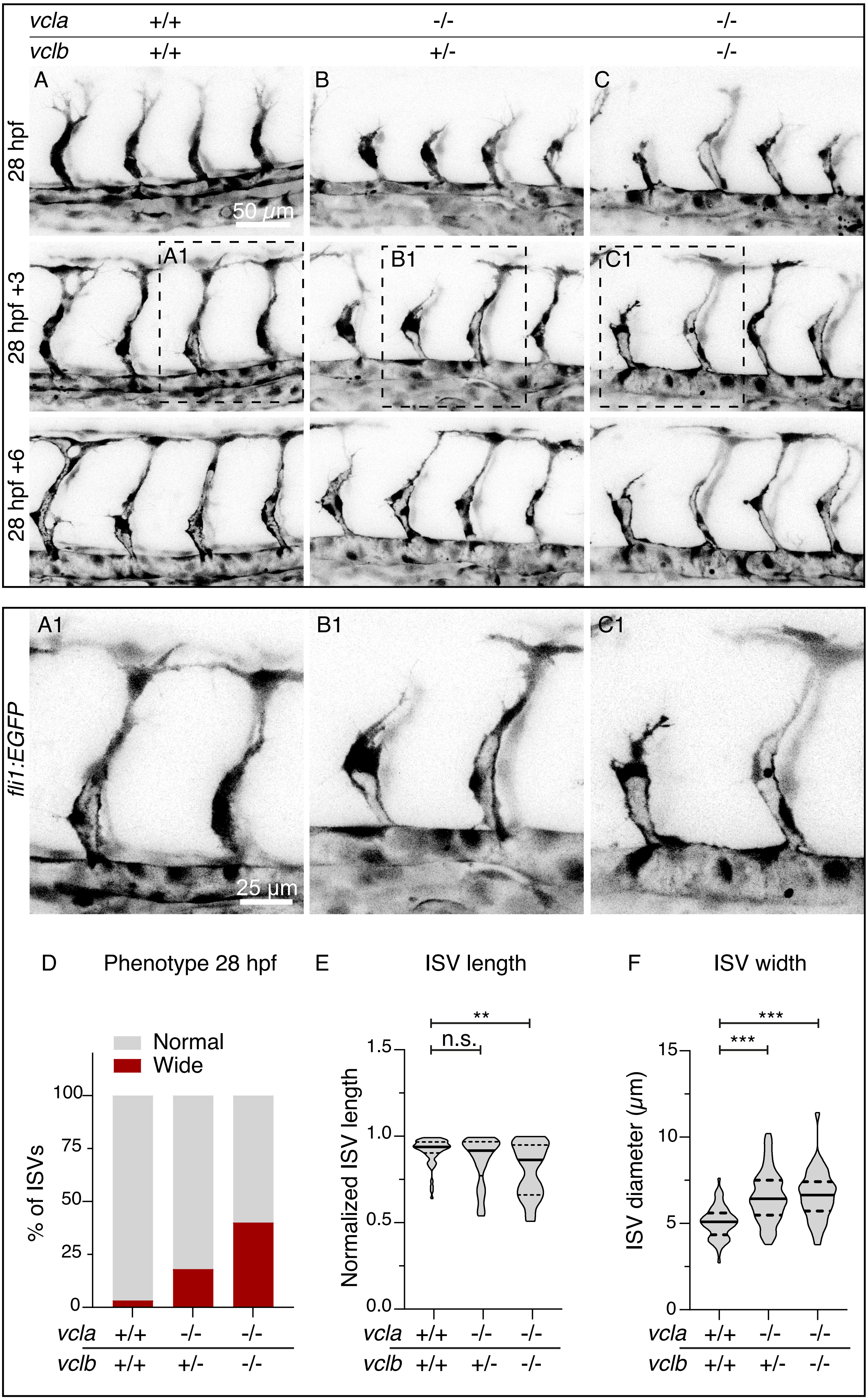


Suppl.Figure3

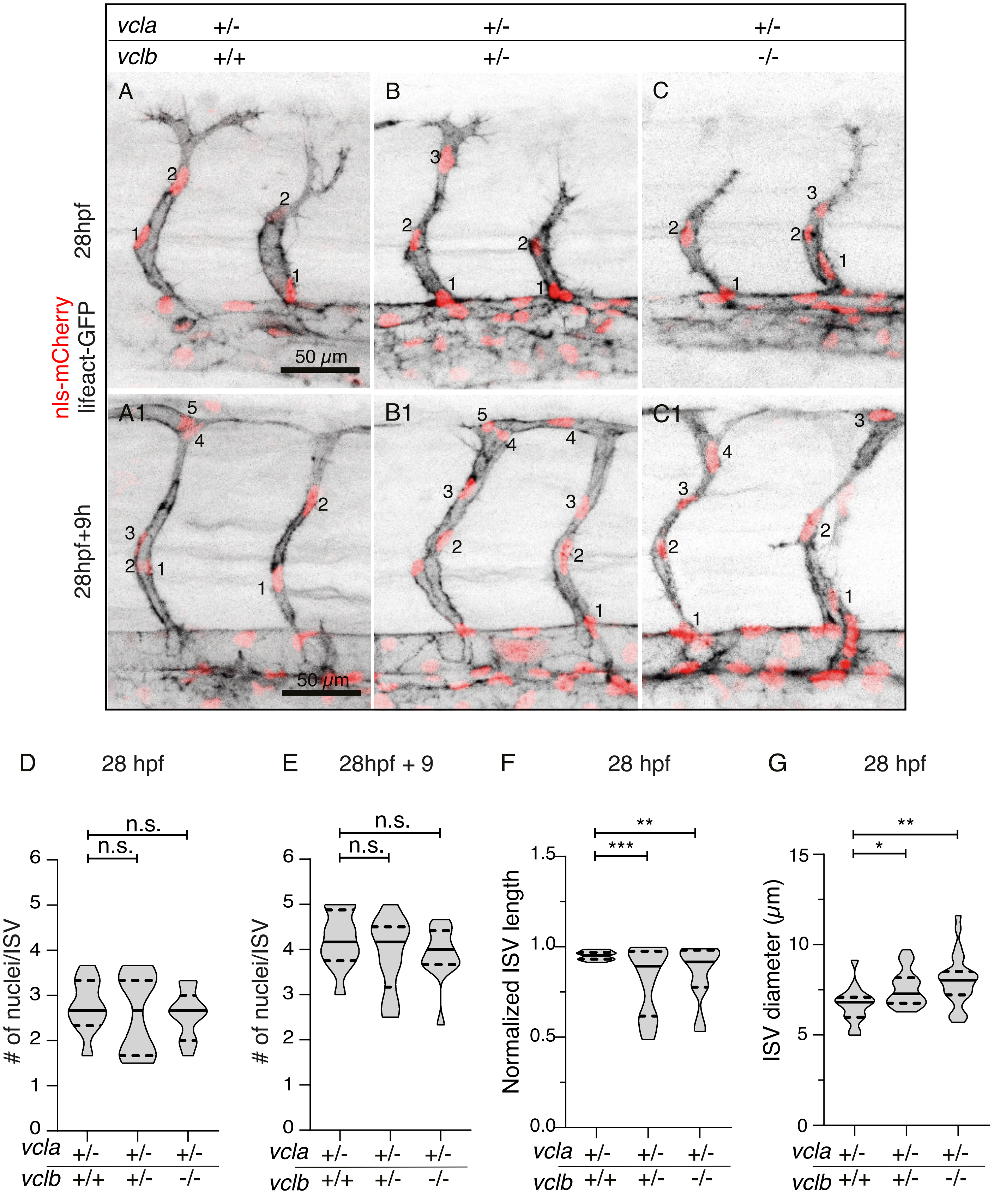


Suppl.Figure4

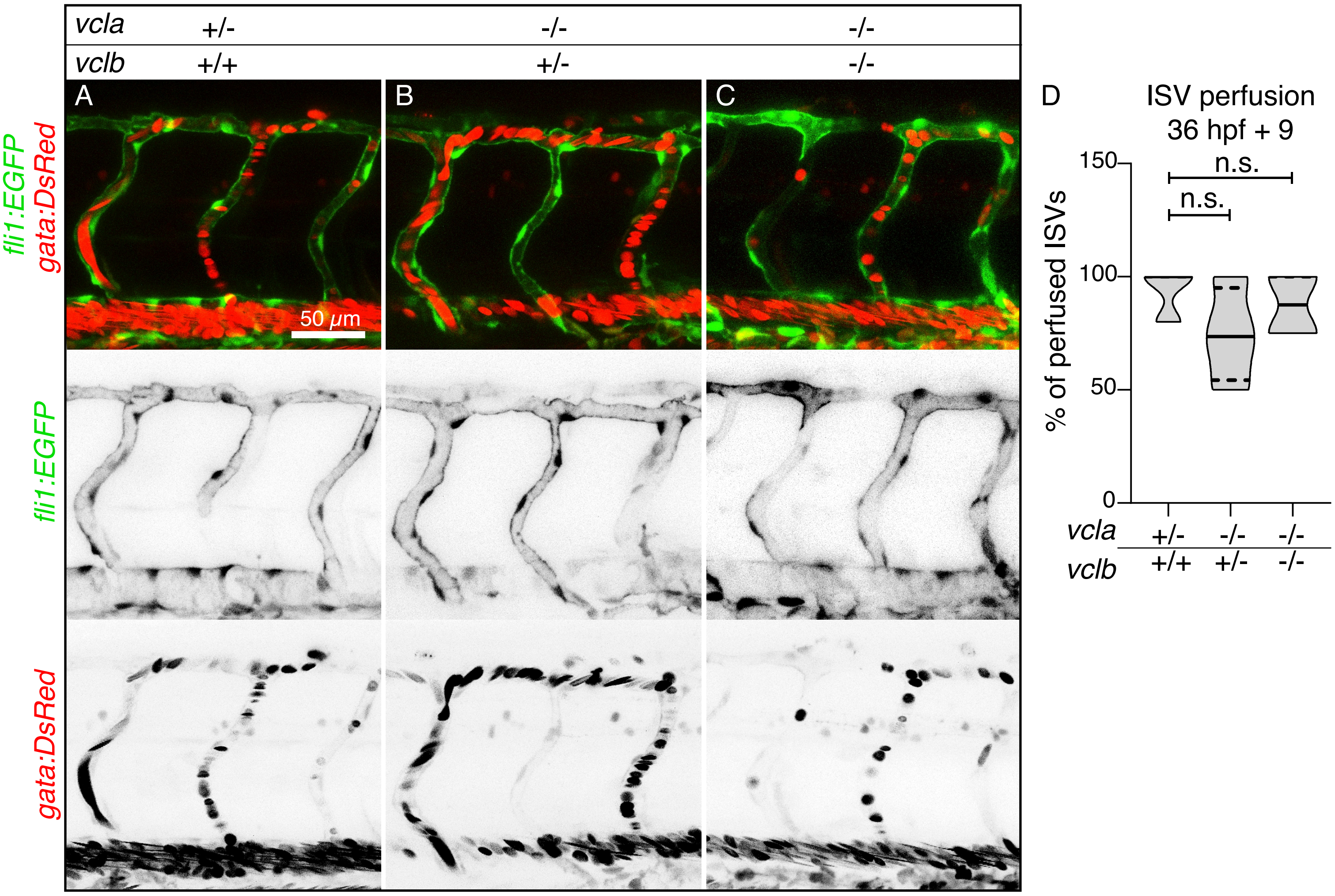


Suppl.Figure5
