## Supplementary material for "Vinculin controls endothelial cell junction dynamics during vascular lumen formation": Supplentary Figure Legends

**Supplemental Figure 1. Hemodynamic forces induce junctional fingers.**

**(A-C)** Stills of time lapse confocal imaging from a representative 36hpf Tg*(kdrl:mCherry-CAAX;fli:pecam1-EGFP*) zebrafish embryo (A) before treatment , (B) after tricaine addition and (C) after norepinephrine addition (110 min) (C). GFP+ signal marks Pecam1 and RFP+ signal indicates the cell membrane (Supplementary movie 3). Scale bar, 10 µm. **(A1-C1)** Inverted images of P1-EGFP of A-C. (A1) Before treatment, junctional fingers are present (black arrows), (B1) while after tricaine treatment junctional fingers rapidly disappear, whereas (C1) sequential treatment with norepinephrine induces finger formation (green arrows). **(A2-C2)** Inverted images of mCherry-CAAX of A-C. (A2) Membrane protrusions (black arrows) are present at the site of junctional fingers before treatment, (B2) which disappear after tricaine treatment. (C2) Treatment with norepinephrine causes reemergence of junctional clusters (green arrows). **(D)** Line graph analysis showing the junctional finger length ±S.D. and lumen diameter over time before (0 to 20 minutes), after treatment with tricaine (25 to 65 minutes) and after treatment with norepinephrine (70 to 100 minutes) from the same *Tg*(*kdrl:mCherry-CAAX;fli:pecam1-EGFP*) embryo.

**Supplemental Figure 2. PCR strategy for *vinculin* isoforms.**

**(A)** Schematic representation of the PCR strategy used to screen for *vcla* mutant embryos or fish. Four allele specific primers can generate different products, depending on the presence of the *vlca* mutation (as described in Han et al, 2017). The specific products generated for the different genotypes are shown in the lower panel. **(B)** Schematic representation of the PCR strategy used to screen for the *vclb* mutant embryos or fish. The combination of four allele specific primers can generate different products, depending on the presence of the *vclb* mutation. The specific products generated for the different genotypes are shown in the lower panel.

**Supplemental Figure 3. Vinculin controls blood vessel morphogenesis during early vascular development.**

**(A-C)** Stills from time lapse imaging of ISVs from Tg*(fli:EGFP)* embryos at 28 hpf with genotypes (A) wild-type (*vcla*^+/+^;*vclb*^+/+^), (B) heterozygous mutant (*vcla*^-/-^;*vclb*^+/-^), or (C) *vcl* double knockout (*vcla*^-/-^;*vclb*^-/-^). Time points 28 hpf, 28 hpf + 3 hours and 28 hpf + 6 hours. Scale bar, 50 µm. **(A1-C1)** Zoom-in images of outlined boxes in A-C. Scale bar, 25 µm **(D)** Bar graph shows the percentage of ISVs with normal (grey) or wide (red) blood vessel phenotype in wild-type (*vcla*^+/+^;*vclb*^+/+^), heterozygous mutant (*vcla*^-/-^;*vclb*^+/-^) or *vcl* double knockout (*vcla*^-/-^;*vclb*^-/-^) embryos. **(E -F)** Violin plot showing the average length of ISV normalized to the maximum ISV length of the embryo (E) and the average ISV diameter (F) of *vinculin* wild-type (*vcla*^+/+^;*vclb*^+/+^) heterozygous (*vcla*^-/-^;*vclb*^+/-^) or *vcl* double knockout (*vcla*^-/-^;*vclb*^-/-^) embryos at 28hpf (n = 12 (*vcla*^+/+^;*vclb*^+/+^), 16 (*vcla^-/-^;vclb*^+/-^) and 8 (*vcla*^-/-^;*vclb*^-/-^), n.s. = non-significant, (E) ** p<0.01 (Kruskal-Wallis test and Dunnett’s post-test), (F) ***p<0.001 (Kruskal-Wallis test and Dunnett’s post-test). **See supplementary movie 5.**

**Supplemental Figure 4. Characterization of the vascular phenotype of *vinculin* mutant embryos.**

**(A-C)** Images of ISVs of *vinculin* single heterozygous (*vcla*^+/-^;*vclb*^+/+^), double heterozygous (*vcla*^+/-^;*vclb*^+/-^) or *vclb* homozygous *(vcla*^+/-^;*vcla*^-/-^) Tg(*fli:lifeact-GFP;kdrl:nls-mCherry*) 28 hpf embryos. **(A1-C1)** Images of ISVs of single heterozygous (*vcla*^+/-^;*vclb*^+/+^), double heterozygous (*vcla*^+/-^;*vclb*^+/-^) or *vclb homozygous* (*vcla*^+/-^;*vcla*^-/-^) embryos Tg(*fli:lifeact-GFP;kdrl:nls-mCherry*) embryos at 28 hpf + 9 hours. Numbers (ie.1,2,3..) indicate endothelial cell nuclei in one ISV. Scale bars, 50 µm. **(D-E)** Violin plot showing the average number of nuclei per ISV in *vinculin* single heterozygous (*vcla*^+/-^;*vclb*^+/+^), double heterozygous (*vcla*^+/-^;*vclb*^+/-^) or *vclb* homozygous (*vcla*^+/-^;*vcla*^-/-^) embryos at respectively 28 hpf (D) and 28 hpf + 9 (E) (n = 10 (*vcla*^+/-^;*vclb*^+/+^), 11 (*vcla*^+/-^;*vclb*^+/-^) and 12 (*vcla*^+/-^;*vclb*^-/-^) embryos) and 28 hpf + 9 hours (E). n = 12 (*vcla*^+/-^;*vclb*^+/+^), 13 (*vcla*^+/-^;*vclb*^+/-^) and 14 (*vcla*^+/-^;*vclb*^-/-^) embryos, n.s. = non-significant (One-way ANOVA and Dunnett’s post-test). **(F-G)** Violin plot showing the average length of ISV normalized to the maximum ISV length of the embryo (F) and the average ISV diameter (G) of *vinculin* wild-type (*vcla*^+/+^;*vclb*^+/+^) *vclb* heterozygous (*vcla*-/-;*vclb*^+/-^) or double homozygous (*vcla*^-/-^;*vclb*^-/-^) 28hpf embryos (n= 5 (*vcla*^+/-^;*vclb*^+/+^), 10 (*vcla^+/-^;vclb*^+/-^) and 8 (*vcla*^+/-^;*vclb*^-/-^)), n.s. = non-significant, ** p<0.01 (Kruskal-Wallis test and Dunnett’s post-test). **See supplementary movie 6.**

**Supplemental Figure 5. Blood flow analysis in *vinculin* mutant embryos.**

**(A – C)** Images of ISVs of *vinculin* single heterozygous (*vcla^+/-^;vclb^+/+^*)*, vclb* heterozygous (*vcla-/-;vclb^+/-^*) *or vcl* double knockout (*vcla^-/-^;vclb^-/-^*) Tg(*fli1:EGFP;gata1a:dsRed*) embryos at 36 hpf + 9 hours . Scale bar, 50 µm. **(A1 – C1)** Inverted images of *fli1:EGFP*. **(A2 – C2)** Inverted images of *gata1a:dsRed*. **(D)** Violin plot showing the percentage of perfused ISVs in single heterozygous (*vcla^+/-^;vclb^+/+^*), *vclb* heterozygous (*vcla^-/-^;vclb^+/-^*) or *vcl* double knockout (*vcla^-/-^;vclb^-/-^*) embryos at 36 hpf + 9 hours, (n = 4 (*vcla^+/-^;vclb^+/+^*), 3 (*vcla^-/-^;vclb^+/-^*) and 2 (*vcla^-/-^;vclb^-/-^*) embryos, n.s. = non-significant (Kruskall-wallis test with Dunnett’s post-test). **See supplementary movie 7.**

**Supplemental movie 1. Junctional finger dynamics during lumen opening.**

Related to figure 2. Time lapse imaging of Tg(*kdrl:mCherry-CAAX;fli:pecam1-EGFP*) embryo starting at 32 hpf with a 24-minute interval.

**Supplemental movie 2. Hemodynamic forces affect junctional finger dynamics.**

Related to figure 4. Zoom in of time lapse confocal imaging of a Tg*(kdrl:mCherry-CAAX;fli:pecam1-EGFP*) 36 hpf embryo before, after treatment with norepinephrine and after treatment with tricaine. 20-minute time interval.

**Supplemental movie 3. Hemodynamic forces induce junctional fingers.**

Related to supplemental figure 1. Zoom in of time lapse confocal imaging of a Tg*(kdrl:mCherry-CAAX;fli:pecam1-EGFP*) 36 hpf embryo before, after treatment with tricaine and after treatment with norepinephrine, 20 minute time interval.

**Supplemental movie 4. Vinculin is recruited to junctional fingers at their maximal size.**

Related to figure 5. Zoom in time lapse imaging of Tg(*fli:pecam1-EGFP*) 36 hpf embryos with transient expression of injected *kdrl:mCherry-vinculin* 20 minute time interval.

**Supplemental movie 5. Vinculin controls blood vessel morphogenesis during early vascular development.**

Related to supplemental figure 3. Time lapse imaging of Tg*(fli:EGFP)* embryos starting at 28 hpf with genotypes wild-type (*vcla*^+/+^;*vclb*^+/+^), heterozygous mutant (*vcla*^-/-^;*vclb*^+/-^), or *vcl* double knockout (*vcla*^-/-^;*vclb*^-/-^). 30 minute time interval.

**Supplemental movie 6. Characterization of the vascular phenotype of *vinculin* mutant embryos.**

Related to supplemental figure 4. Time lapse imaging of Tg(*fli:lifeact-GFP;kdrl:nls-mCherry*) embryos with genotypes: single heterozygous (*vcla*^+/-^;*vclb*^+/+^), double heterozygous (*vcla*^+/-^;*vclb*+/-) or *vclb* homozygous (*vcla*^+/-^;*vcla*^-/-^) starting at 28 hpf with a 30 minute interval.

**Supplemental movie 7. *vinculin* mutants do not show alterations in blood flow.**

Related to supplemental figure 5. Time lapse imaging of with genotypes single heterozygous (*vcla^+/-^;vclb^+/+^*)*,* heterozygous (*vcla^-/-^;vclb^+/-^*) *or vcl dko* (*vcla^-/-^;vclb^-/-^*) Tg(*fli1:EGFP;gata1a:dsRed*) embryos. Starting at 36 hpf with a 30-minute time interval.
